# A comprehensive study of the microbiome and resistome of chicken waste from intensive farms

**DOI:** 10.1101/2022.07.04.498670

**Authors:** Aleksandra Błażejewska, Magdalena Zalewska, Anna Grudniak, Magdalena Popowska

## Abstract

The application of chicken waste to farmland could be detrimental to public health. It may contribute to the dissemination of antibiotic resistance genes (ARG) and antibiotic-resistant bacteria (ARB) from feces and their subsequent entry to the food chain. The present study analyzes the metagenome and resistome of chicken manure and litter obtained from a commercial chicken farm in Poland. ARB were isolated, identified and screened for antibiogram fingerprints using standard microbiological and molecular methods. The physicochemical properties of the chicken waste were also determined. ARG, integrons, and mobile genetic elements (MGE) in chicken waste were analyzed by high-throughput SmartChip qPCR. The results confirm the presence of many ARGs, probably located in MGE, which can be transferred to other bacteria. Potentially pathogenic or opportunistic microorganisms and phytopathogens were isolated. More than 50% of the isolated strains were classified as multi-drug resistant, and the remainder were resistant to at least one antibiotic class; these pose a real risk of entering groundwater and contaminating the surrounding environment. Our results indicate that while chicken manure can be sufficient sources of the nutrients essential for plant growth, its microbiological aspects make this material highly dangerous to the environment.

## 1. Introduction

Chicken meat production is considered one of the fastest-growing sectors of the Polish food industry. The US Department of Agriculture (USDA) reports that Poland is one of the largest poultry meat producers in the European Union (EU), with chicken production reaching 2.2 million metric tons in 2020 (USDA, 2020). Total chicken meat production in the EU peaked in 2018, reaching 15.2 million tons; this year, Poland produced almost 17% of this value (176.7 mln chickens, including 192.1 mln laying hens), giving it a leading position within the European Union (Eurostat) [1].

A number of public and environmental health organizations, such as the World Health Organization (WHO), have raised concerns about the spread of antimicrobial resistance (AMR) in bacteria, driven by the extensive use of antibiotics in poultry and livestock production [2]. The most commonly-used antimicrobials in the poultry industry are polymyxins, penicillins, tetracyclines, and trimethoprim with sulfonamides. Antimicrobials like fluoroquinolones, third- and fourth-generation cephalosporins, macrolides, glycopeptides, and polymyxins are considered by the WHO as the “highest priority critically-important antibiotics for human medicine” due to the limited availability of alternatives for the treatment of bacterial infections. While many of these are approved for use in poultry in the largest poultry-producing countries, fluoroquinolones are banned in the United States and cephalosporins in the EU [3].

Antibiotics are administered to treat intestinal infections such as colibacillosis, necrotic enteritis, and other diseases generally caused by *Salmonella* spp., *Escherichia coli*, or *Clostridium* spp. These infections are a major concern among poultry producers, leading to enormous economic losses [3]. However, up to 70% of the administered antimicrobial compound is excreted with feces and urine in their active form and remains in fertilized soil for a prolonged time [4,5]. Furthermore, antibiotics are generally administered to the entire flock, not only to isolated infected birds, with the aim of treating disease (therapy), preventing infection (metaphylaxis), and promoting growth [6]. The birds are, in most cases, kept in high numbers in intensive breeding facilities and are often transported at high densities over long distances, which makes them especially prone to disease. The type and extent of antibiotics vary between countries based on their economy, level of development, intensity of animal husbandry and animal species [7]. In addition, the method of administration and the volume of antibiotics used vary according to the stage of production and risk of disease [8].

Antibiotic residues are considered an important environmental contaminant. They are believed to play a key role in enriching soil microbiomes with antibiotic-resistant bacteria (ARB) and antibiotic resistance genes (ARG) by creating on-site pressure, which facilitates the development of new ARG and supports the transfer of existing ones between bacteria [9–11].

In Poland, 86.8% of chicken is raised in the cage bedding system (source of manure), 9.6% in the bedding system (source of chicken litter), 3.2% in the free-range system, and 0.3% in the organic system. A similar trend can be seen across Europe, with cage bedding systems dominating (58.7%); however, a higher percentage of chickens are raised in the bedding system (27.5%), free-range system (9.2%), and organic system (4.6%) [12]. While farms caged layers and broiler breeders produce chicken manure consisting of only fecal droppings (undigested food, urea, gastrointestinal microbiota), the litter from floor-raised chickens consists primarily of animal droppings (feces with gastrointestinal microbiota) mixed with bedding material, such as sawdust, and smaller amounts of feathers and dropped fodder [13]. Some studies have shown that chicken litter contains a complex and dynamic microbiota composed primarily of gastrointestinal tract and environmental bacteria, depending on the litter management regimens [14]. It is estimated that 1000 broilers can produce approximately 120 kg of feces per year [15]. In Poland, the total manure produced by chicken farms in 2017 was estimated to be approximately 4,494,639 tones; hence, the storage and utilization of chicken waste can present a huge challenge for the whole industry. In developing countries, waste is most commonly managed by spreading unprocessed manure and litter on the soil, with or without prior dilution. Indeed, such natural fertilizer application can significantly improve soil properties and fertility, as chicken waste is rich in nitrogen and contains high quantities of phosphorus and potassium, essential for plant growth [12].

However, despite the clear advantages of animal waste application, this practice has unavoidable risks. The contamination of the soil microbiome, one of the richest and most diverse environments, with antibiotics and antibiotic residues may contribute to the spread of antimicrobial resistance [13], with the antibiotics, growth hormones and potential pathogens living in the chicken intestine playing a significant role [16]. Usually, chicken waste is inhabited by diverse microbes, especially Gram-negative bacteria, some of which are potential human pathogens (*E. coli, Mycobacterium* spp., *Salmonella* spp.). As these microbes can harbor high levels of ARG and ARB, the widespread application of untreated waste in the field can present a hazard to the environment [17]. Chicken waste has also been identified as a major reservoir of mobile genetic elements (MGE), facilitating the exchange of ARG between bacteria via horizontal gene transfer (HGT) [18]. When resistance genes are located on plasmids, transposons or integrons can spread among even non-related bacteria [10]. Recent studies report that phylogeny is one of the main drivers for transferring mobile ARG in animal and human gut microbiota, and that these elements can also be harbored by human pathogens [19].

Fertilizer consisting of chicken manure is characterized by a relatively high prevalence of ARG, reaching up to 60% of isolated bacteria. More importantly, most of these are classified as multi-drug resistant (MDR), being resistant to more than three classes of antimicrobials [20]. There is hence a need for novel legal regulations regarding animal waste treatment, and this requires a clearer understanding the impact of the field application of chicken waste on soil ARB and ARG levels, and their dissemination. Such field application may significantly impact the soil resistome, altering natural soil microbial communities.

The aim of the present study was to evaluate the potential of chicken waste (manure, litter) as a source of hazardous contaminants released into the natural environment, and as a crucial hot spot for the spread of antimicrobial resistance. Currently, little data exists on the distribution of ARGs and ARB in chicken manure, especially that derived from Poland, one of the main chicken meat producers in the EU. To assess the potential threat, the study analyzed the microbial community composition of two types of chicken waste, viz. chicken manure (CM) and chicken litter (CL), based on sequencing protocols targeting the V3-V4 variable regions of 16S rRNA. The resistome of samples was analyzed by high-throughput qPCR, targeting 384 genes in each sample, to confirm the presence of ARG. The study also attempted to isolate selected antibiotic-resistant, potentially pathogenic bacteria listed by the WHO as the so-called ‘priority pathogens list for R&D of new antibiotics’ from the CL and CM [21].

## 2. Results and discussion

### 2.1. Physicochemical characterization of chicken wastes

The data on the physicochemical properties of the two kinds of tested chicken waste are presented in Table 1. Typical arable soils found in Poland (sandy, with low humus content, soil evaluation class IV, non-fertilized) were also analyzed as a reference. The collected data shows that the chicken manure (CM) was slightly alkaline (pH=8), while the litter (CL) samples had a closer pH to the soil value (pH=6.5); i.e. more neutral and closer to healthy and fertile soil. The pH value of our CL samples differs from those obtained by Dumas et al. [22], but pH value is inseparably related to the moisture content, and this may differ between sample sites; for example, one may be closer to a water source. Research from 2010 indicates that the application of chicken manure can significantly increase soil pH [23]. Both manure samples have higher levels of macronutrients such as magnesium (Mg), calcium (Ca), and phosphorus (P) and heavy metals like copper (Cu) and zinc (Zn) compared to soil. Even so, the heavy metal concentrations in the collected samples were below the maximum permissible limits established by European Parliament and the Council in the Regulation (EC) No 1069/2009 from October 21, 2009 [24]. Per kilogram of dry matter, organic fertilizer may contain a maximum of 5 mg Cd, 2 mg Hg, 100 mg Cr, 60 mg Ni, and 140 mg Pb. CL was almost three times richer in organic matter (OM; 60.2%) and two times richer in total nitrogen (2.78%; %wt N) than CM, with only 18.4% OM and 1.22%wt N. At the same time, the CM samples had more ammoniacal nitrogen (3200 mg/kg), than the CL sample with 2400 mg/kg (Table 1). This disproportion may be due to the differences in the composition of the analyzed waste types: chicken litter additionally has bedding material [13].

**Table 1.**
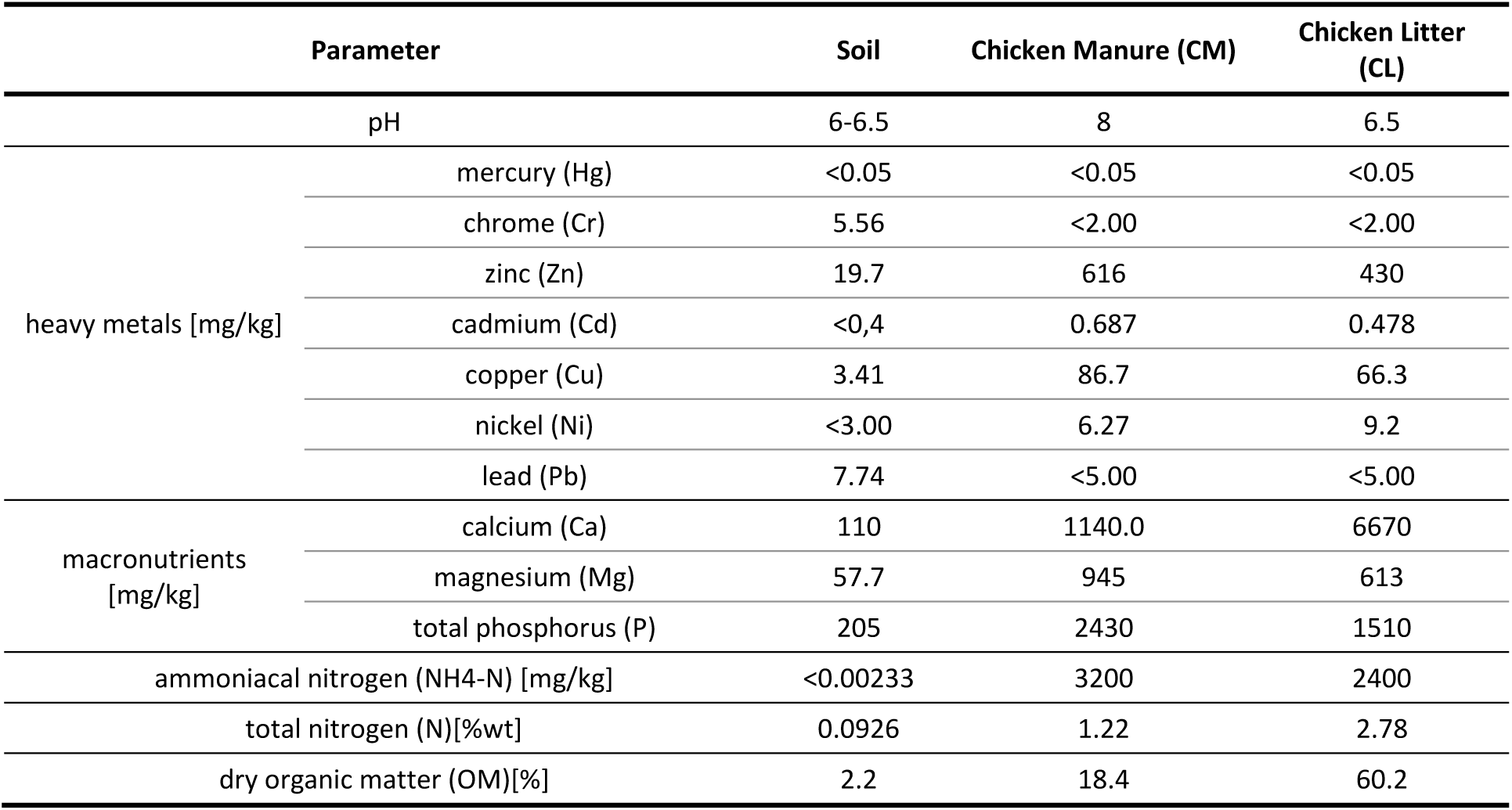
Physicochemical properties of the two kinds of chicken manure from Polish commercial farm and non-manured soil.

It is hard to compare the physicochemical characteristics of environmental samples with other those of studies due to the lack of unified methods and sample parameters. Even so, our results, especially those of the CM samples, seem to be comparable to those of other samples collected from similar sites in other big poultry-producing countries. In poultry waste collected from several sampling sites in Southern Brazil (one of the biggest poultry meat exporters), the average dry matter content was 64.3%, total nitrogen content was 2.2 % (similar to the value of our CL samples) and the pH was also alkaline: most samples were higher than pH 7, which is slightly more alkaline than the CL samples collected for our experiments [25]. In addition, previous studies on composting chicken manure in Manjung Region, Malaysia, found a higher total nitrogen content (5.52%) and a much lower pH (6.1) than our CM samples [26]. Our results were similar to those of chicken manure samples from South Africa, with a pH of 7.97 to 6.94 and total nitrogen content ranging from 1.6 - 3.2% depending on the sampling site. In addition, the total phosphorus content in collected samples was also similar to our results, ranging from 1963 mg/kg to 2644 mg/kg. The high phosphorus levels in poultry manure and litter occur mainly because chickens only utilize a minimal portion of the supplied phosphorus, with the rest being excreted with feces or urine [27].

Our physiochemical analysis also revealed high levels of calcium (Ca) and phosphorus (P) in the chicken feces. Calcium (Ca) and phosphorus (P) are crucial nutrients: they are linked to many biological processes, such as cell proliferation, bone formation, blood clotting and energy metabolism, and in chickens, disorders in Ca and P homeostasis are associated with a decline in growth and egg-laying performance. The main sources of Ca and P for laying hens are mineral supplements and plant-derived compounds. In addition, to meet the required levels, a highly-productive laying hen diet is supplemented with high-quality inorganic phosphates. However, due to their inefficient use of P, livestock and poultry are significant P producers and, thus, a major source of P input into the environment. The mechanisms of P homeostasis are strongly conserved and linked to Ca metabolism: the dietary Ca/P ratio has a strong impact on health and performance, and so must lie within a physiological range [28]. Imbalances in dietary Ca and P can also result in excess P excretion, which can have negative environmental effects when poultry litter is applied as a fertilizer to the soil, causing eutrophication and environmental pollution [29]. Due to the importance of phosphorus and calcium for chicken growth and performance, chicken diets are typically overloaded with these two minerals to reduce the likelihood of deficiency [30].

In addition, a higher level of calcium was noted in feces from broiler chickens (CL) than from laying hens (CM). Typically, laying hens are additionally supplemented with calcium to provide for eggshell formation and yolk production during the laying period: calcium accounts for 40% of the eggshell weight in the form of CaCO3 [28]. Our results suggest that the broiler chickens and laying hens used in the study may get similar amounts of calcium in their regular diet, but laying hens use more calcium during every-day performance. However, excess levels of dietary Ca can reduce the performance of chickens and inhibit growth, and reduce weight gain and feed efficiency [31].

### 2.2. Bacterial community composition

#### 2.2.1. Identification of bacteria species and antibiotic susceptibility testing

Initial isolation of bacteria species using selective media supplemented with antibiotics resulted in the collection of 649 strains. After eliminating potentially repetitive strains, about 119 we obtained for final identification and antimicrobial susceptibility analysis. The differences in bacteria species composition between the two analyzed types of chicken waste (CM and CL) are shown in Figure 1. A greater diversity of bacterial strains were isolated from CL than those from CM. In the CM, only Gram-negative, opportunistic pathogens such as *Myroides odoratus* (1.7%), *Providencia rettgeri* (2.5%), and *Ochrobactrum intermedium*

**Figure 1.**
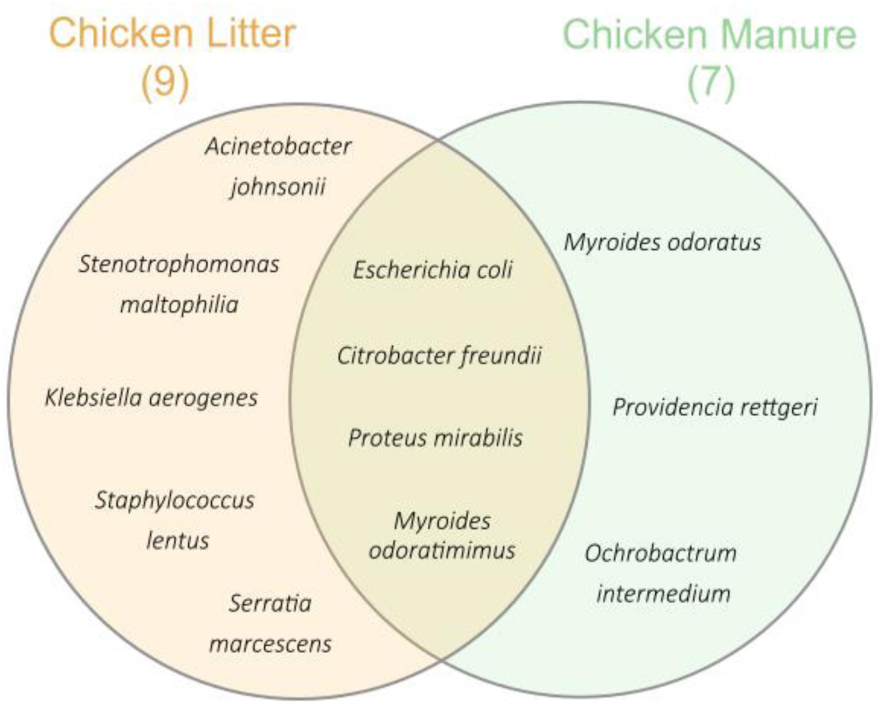
Venn Diagram presenting the differences in bacteria species found in two analyzed types of chicken waste – CM and CL

(0.8%) were isolated. Species like *Citrobacter freundii* (0.8%), *Proteus mirabilis* (23%), *E. coli* (19%), and *Myroides odoratimimus* (23.5%) were present in both waste types. The CL waste was dominated by Gram-negative bacteria species, such as *Acinetobacter johnsonii* (0.8%), *Klebsiella aerogenes* (2.5%), *Stenotrophomonas maltophilia* (0.8%) and *Serratia marcescens* (3%); however, one Gram-positive bacterium was isolated: *Staphylococcus lentus* (16%).

Depending on the identified species, different AST cards were used to test susceptibility to groups of antibiotics. The percentage of isolates that are sensitive, intermediate, or resistant to the selected antibiotics is shown in Table S1. Among 91 tested strains, 61.5% were resistant to ciprofloxacin, 50.5% gentamicin, and 25.6% trimethoprim/sulfamethoxazole.

Out of 72 tested strains, 16.7% were susceptible to piperacillin/tazobactam, 7% to imipenem, 2.5% to meropenem, 18.1% to amikacin, 54.2% to tobramycin and 36.4% to tigecycline. In addition, 57 strains were tested for susceptibility to second, third and fourth generation cephalosporins: 36.8% were resistant to cefuroxime, 52.6% to ceftazidime, 48.3% to cefotaxime, 1.7% to cefepime, 54.4% to cefuroxime-axetil. The Gram-positive strains (19 strains of *Staphylococcus lentus*) were tested against three common antibiotics: 47.9% of the strains were resistant to erythromycin, 79% to clindamycin, and all were resistant to tetracycline (Table S1).

From both of waste types, 23 strains of *E. coli*, 13 strains of *M. odoratimimus*, and one strain of *C. freundii* were isolated and identified. All were resistant to more than five anti-biotics. From CL, five strains of *S. marcescens* were found to be resistant to at least three antibiotics, three strains of *K. aerogenes* resistant to amoxicillin/clavulanic acid and cefuroxime, and one strain of *A. johnsonii* resistant to piperacillin/tazobactam, ciprofloxacin, and piperacillin. From CM, three *P. rettgeri* strains and one *M. odoratus* strain was found to be resistant to six or more antibiotics. The complete phenotypes of the isolated and identified strains are presented in Table 2.

**Table 2.**
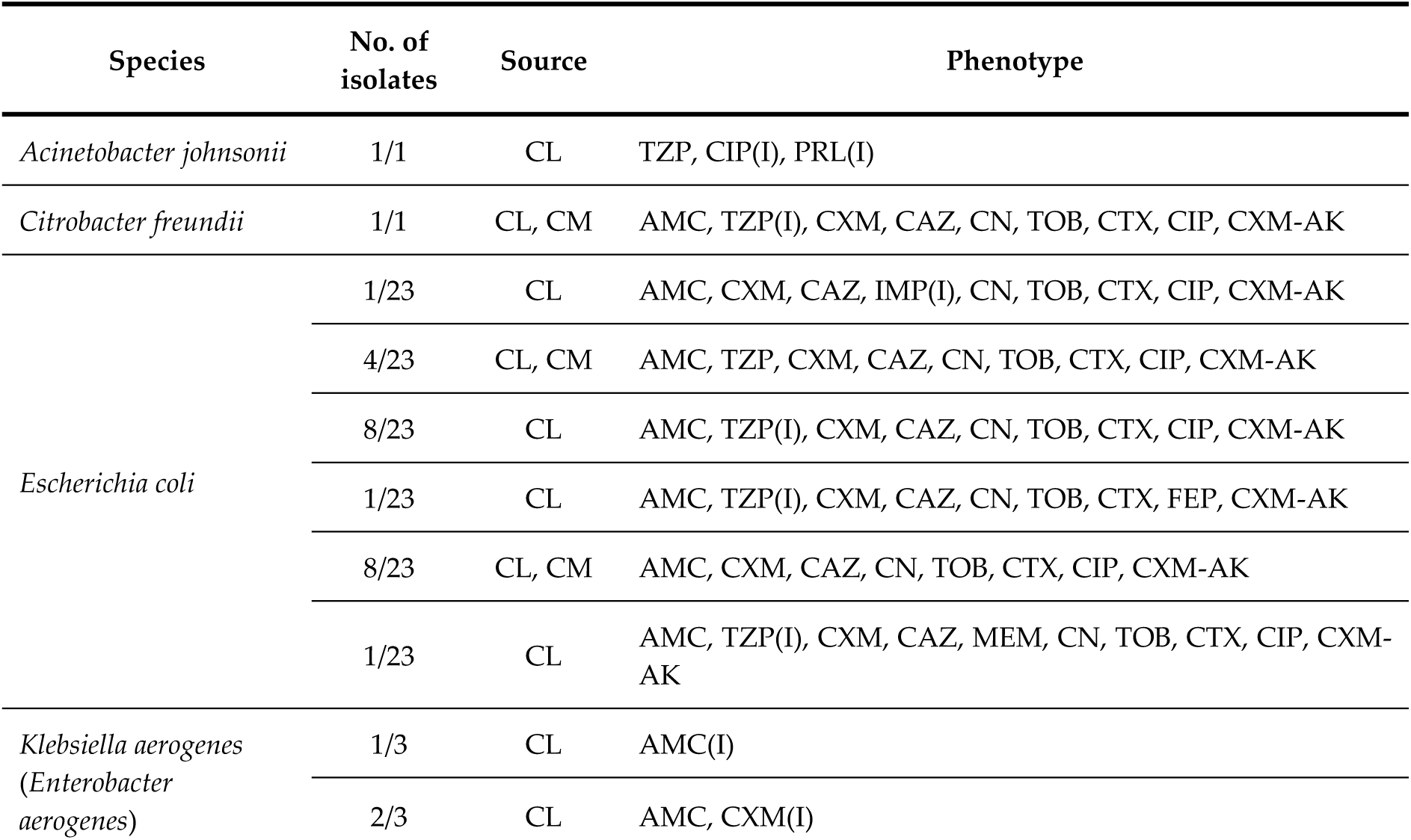
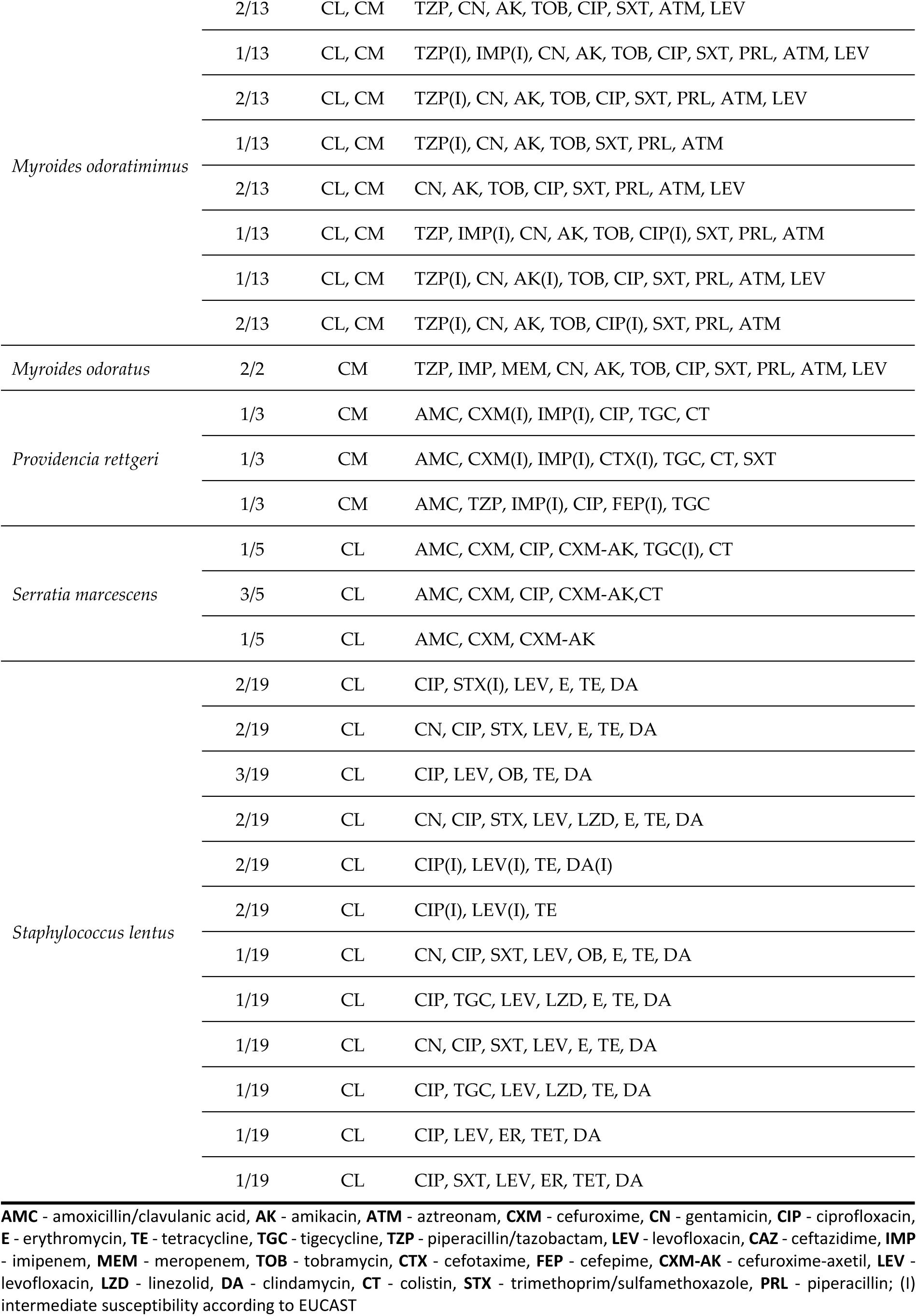
Antibiotic resistance phenotypes of isolated strains form two types of chicken wastes – CL and CM

Among the samples, *M. odoratimimus* was the most prevalent of the isolated bacteria. *Myroides* spp. are common residents in natural environments like water and soil. However, these Gram-negative bacilli can carry resistance genes to multiple antibiotics and, as opportunistic pathogens, may cause infection in humans, especially in immunosuppressed patients [32]. In 2010, an outbreak of *Myroides* spp. caused several urinary tract infections in Tunisian hospitals. The disc diffusion susceptibility test showed that all isolates were resistant to the β-lactam antibiotic imipenem [33]. Our isolates also showed intermediate resistance to imipenem (intermediate being defined according to EUCAST recommendations ISO 20776-1). Additionally, our isolates were resistant to ciprofloxacin, gentamicin or tobramycin.

*E. coli* is another species commonly present in chicken wastes. This coliform is one of the main residents of the chicken intestine and can survive in natural environments like water and soil [34]. It is also considered a marker of environmental contamination with fecal matter [35]. In research from Cameroon, the overall percentage of resistant *E. coli* isolates from the chicken litter was approximately 58.4%. After susceptibility testing, almost 83% of them were resistant to more than three antibiotics and considered MDR strains. Similar findings were obtained for the phenotypes of the *E. coli* isolates in our study, confirming a high level of MDR among the species, with all 23 isolated *E. coli* being resistant to at least eight antibiotics. In the samples collected in Cameroon, the *E. coli* demonstrated lower levels of resistance to ciprofloxacin (36%) and imipenem (45%) than to ampicillin (91%) and amoxicillin with clavulanic acid (89%) [36]. In our isolated *E. coli* strains, resistance to ciprofloxacin (61.5%) was similar to that of amoxicillin (63.2%), with resistance to imipenem at a much lower level (7% of isolated bacteria). Furthermore, in *E. coli* strains isolated from organic poultry fertilizer in Brazil, most were resistant to more than five tested antimicrobials [37].

Moraru et al. [38] report the presence of fluoroquinolone- and ciprofloxacin-resistant *Enterobacteriaceae* in chicken manure. During our study, the most common form of resistance among the isolated strains (n=91) was to ciprofloxacin (CIP) (61.5%) (Table S1). This broad-spectrum fluoroquinolone is commonly used against Gram-negative bacteria in veterinary contexts to prevent avian colibacillosis.

These findings are significant, especially from the environmental point of view, as chicken litter is commonly used as natural fertilizer on arable fields. These practices carry a high risk of potential crop contamination; indeed, a case study on gardening crops from Southern Benin confirmed crop contamination by fecal coliforms including *E. coli* after poultry manure fertilization [39]. Moreover, bacteria from *Providencia* spp., considered commensals in the gastrointestinal tract, pose a threat to humans and animals as opportunistic photogenes and can cause severe gastric infections. In our studies, three isolates of *P. rettgeri* were resistant to at least three tested groups of antibiotics (beta-lactams, fluoroquinolones, sulfonamides, and polymyxins), which allows them to be classified as possessing an MDR phenotype. In total, 19 Gram-positive *S. lentus* were isolated in the present study, only from CL, and all were resistant to at least three antibiotic groups. Graham et al. [40] report that some staphylococci carrying resistance to more than three antibiotics are able to survive even a 120-day storage period in a two-walled, roofed shed. The possession of an MDR phenotype makes any bacterium even more dangerous: not only can it limit potential treatment strategies in case of infection, environmental bacteria may transfer resistance vertically or horizontally to other bacteria [10].

In many countries, e.g., the US, chicken litter is commonly used as a bedding material for young broiler chicks placed in chicken houses. After being placed in commercial chicken houses where litter serves as the bedding material, chicks are exposed to bacteria that can enter the immature gut. As the gastrointestinal tract of young chickens offers weak resistance to colonization, it can be easily colonized by bacteria. Beginning from day one, chicks begin pecking at and consuming litter materials, inoculating their young gastrointestinal tract with bacteria present in the litter. Thus, the litter microbiome may affect the development process of intestinal microbiota and its eventual composition and structure in chickens [13]. Our study showed that chicken litter carries many potentially pathogenic bacteria that are resistant to commonly-used antibiotics, and which may be transferred to young chicks via bedding materials. The colonization of poultry gastrointestinal tract with ARB can cause the further spread of resistance among other chickens and eventually enter the human food chain with contaminated meat.

#### 2.2.2. Microbial community composition

The microbial community composition of the CM and CL samples was determined by targeted sequencing of the variable V3-V4 region of the 16S rRNA gene, and their richness and diversity were analyzed. Among all tested samples, a total number of 21 phyla were identified. In chicken manure, *Bacteroidetes* (41%) were considered the dominant bacterial phyla followed by *Firmicutes* (17%), *Proteobacteria* (12%), and *Spirochaetes* (11%). In CL, the dominant phyla were *Firmicutes* (43%), followed by *Bacteroidetes* (28%), *Proteobacteria* (19%), and *Actinobacteria* (9%). Microbiome compositions for both types of feces are presented as relative abundance bar plots (Figure 2).

**Figure 2.**
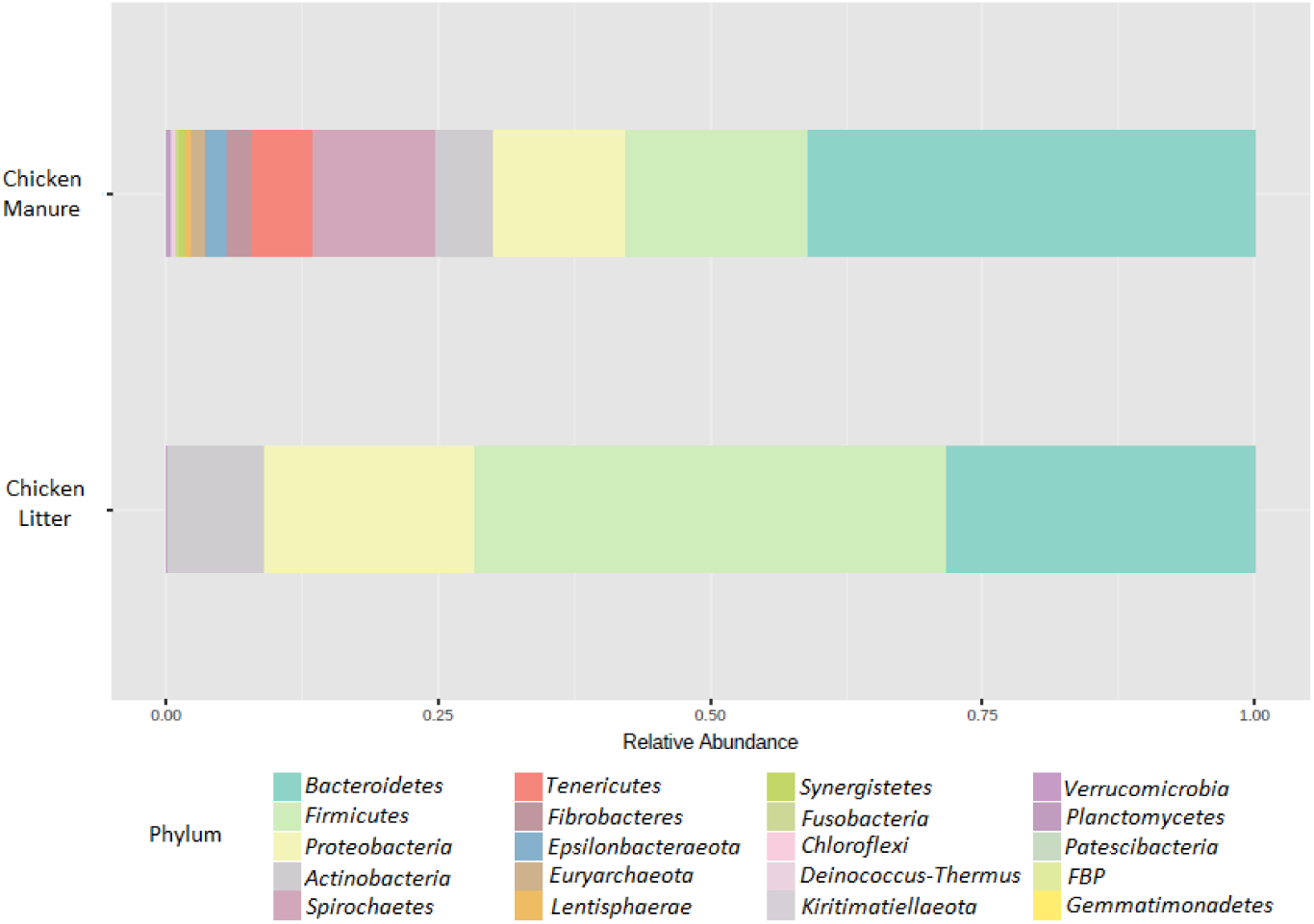
Microbial composition of collected feces samples at the phylum level. The visualization represents the mean value for three replicates.

One of the dominant phyla in our samples was *Bacteroidetes*. This group is a very diverse bacterial phylum that can colonize various environments, including soil, water and, most importantly, the gastrointestinal tract. *Bacteroidetes* act as symbionts in the gastrointestinal tract, mainly helping degrade biopolymers such as polysaccharides. Despite their beneficial contribution to the environment, some members of this phylum are well-known opportunistic or clinically-relevant human and animal pathogens (*Myroides* spp., *Sphingobacterium* spp.) or phytopathogens, like *Flavobacterium johnsoniae* [41]. Due to their importance in depolymerizing or degrading organic substances, their abundance in ecosystems might enhance the rate of organic matter turnover, leading to enhanced CO2 emissions [42]. Its abundance also correlates strongly with pH, with *Bacteroidetes* being more abundant In environments with more alkaline pH values (approximately pH 8), than in those with more neutral pH values (approximately pH 6.5-7) [43]. This observation was also confirmed in our samples, in which *Bacteroidetes* were more abundant in CM (pH 8) than in CL (pH 6.5).

The most abundant phylum in the CL samples was *Firmicutes*. This observation agrees with those of Borda-Molina et al. [44], which identify *Firmicutes* as the most abundant bacterial phylum in the chicken gastrointestinal tract. However, our results for the CL differ from those presented by Gurmessa et al. [42], who report the contents to be *Firmicutes* (54.83%), *Actinobacteria* (33.00%), *Bacteriodetes* (6.86%) and *Proteobacteria* (2.90%); however, like Gurmessa et al [42], the three most abundant phyla constituted approximately 90% of the overall bacterial community composition. Other studies have found *Firmicutes* and *Bacteroidetes* to represent almost 90% of healthy gut microbiota in chickens [45]. *Firmicutes* are also responsible for the degradation of polysaccharides and butyrate production in the chicken gastrointestinal tract. Previous taxonomic analyses based on 16S rRNA sequence assignment by Zhang et al. [46] identified 16 phyla, with the four most dominant being similar to our present findings. It appears that chicken manure samples are, in general, mainly dominated by these four phyla (*Bacterioidetes, Firmicutes, Proteobacteria, Actinobacteria*). In another study, fresh manure and composted samples were dominated by *Actinobacteria* (43.3 vs. 32.6%), *Firmicutes* (38.4 vs. 39.1%), *Proteobacteria* (10.4 vs. 13.5%), and *Bacteroidetes* (6.4 vs. 7.5%) [47]. Ziganshin et al. [48] found *Firmicutes* (23–79%) and *Bacteroidetes* (8– 44%) to be the most abundant phyla in raw chicken manure. However, environmental conditions have been found to vary between poultry houses, and these differences can have a major effect on microbial dynamics, even at the microscale level; spatial scaling of microbial diversity can exist even on a scale of a few centimeters [14].

#### 2.2.3. Microbial diversity

The microbial richness and diversity of the tested samples were analyzed with the Chao1 (Figure 3A) and Shannon (Figure 3B) indexes. The results for the groups were compared using the nonparametric Mann-Whitney test. Higher diversity (*p*≤0.1) and richness (*p*≤0.1) values were observed for CM samples than for CL samples, but only as a general trend. In addition, the beta diversity analysis identified some differences in microbial community structure between the two types of chicken waste (*p*≤0.1), but also only at the trend level; the results were visualized by PCoA (Principal Coordinate Analysis) based on the Bray-Curtis dissimilarity matrices (Figure 4).

**Figure 3A.**
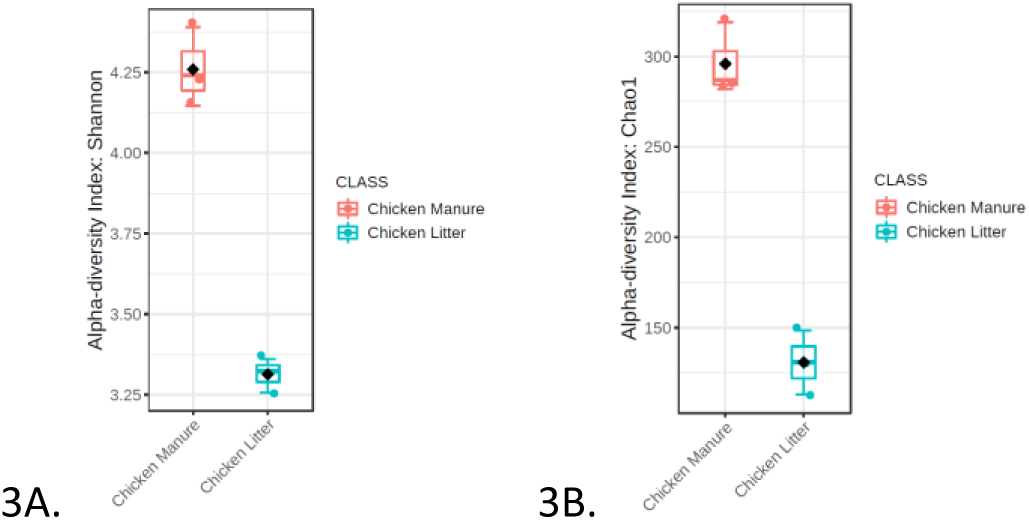
Shannon indexes of microbial community diversity *p*≤ 0.1; statistical analyses were performed with the Mann-Whitney test **Figure 3B**. Chao 1 indexes of microbial community richness *p*≤0.1; statistical analysis was performed with the Mann-Whitney test in collected chicken waste samples (Chicken Manure: CM_1, CM_2,CM_3 [orange dots] and Chicken litter: CL_1, CL_2, CL3 [blue dots]).

**Figure 4.**
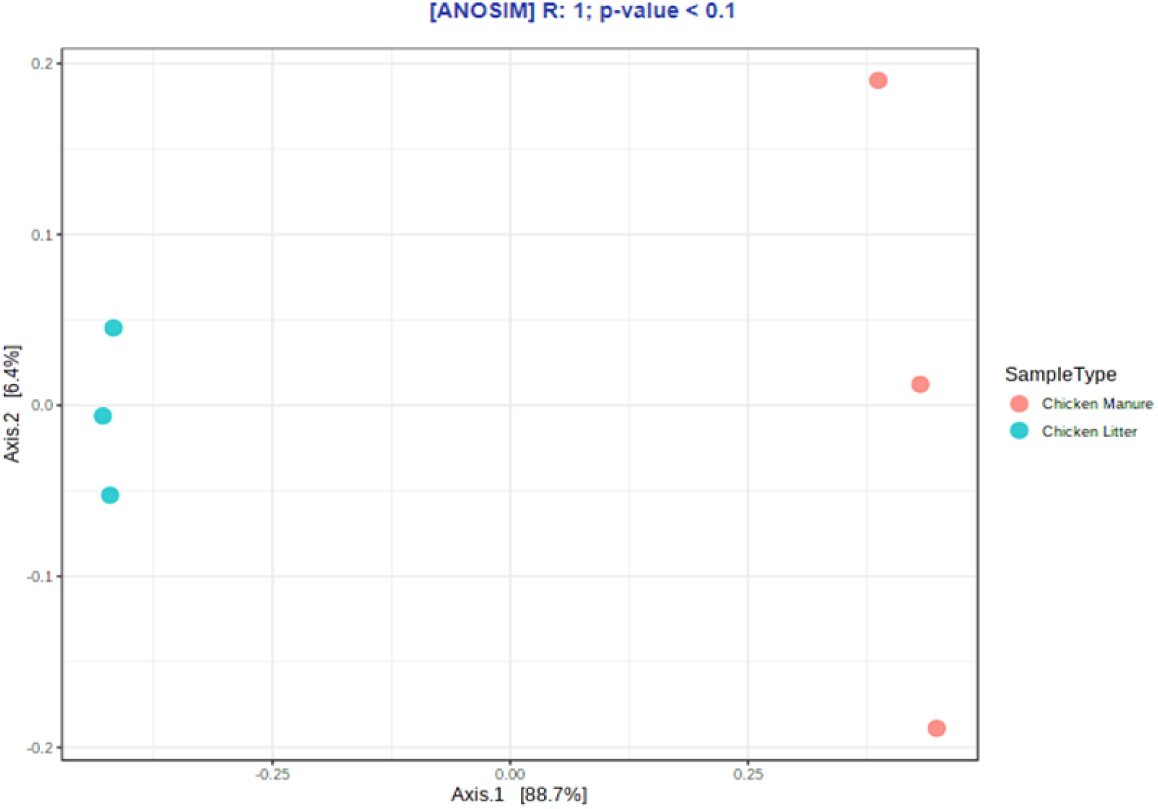
Principal coordinate analysis (PCoA) of microbial community compositions based on the Bray-Curtis dissimilarity matrices [ANOSIM] in collected chicken waste samples (Chicken Manure: CM_1, CM_2,CM_3 [orange dots] and Chicken litter: CL_1, CL_2, CL_3 [blue dots]).

Considerable differences in diversity and richness were observed in the collected samples, and this may be due to the fact that α-diversity is strongly influenced by the physicochemical conditions of the waste and environmental factors. Few studies have compared the antimicrobial communities of wet (manure) and dry (litter) chicken waste. The fact that chicken litter is a mixture of poultry manure and bedding likely influences the physiochemical characteristics of the resulting litter, which subsequently affects the bacterial community structure [42]. Our present findings indicate that CL and CM differ most prominently with regard to moisture, pH value, total nitrogen concentration and dry organic matter content. All these characteristics can affect the richness and diversity of the bacterial community, which in turn are correlated with the moisture level of the sample: in CM, a higher moisture content can increase α-diversity by enhancing the transport of dissolved nutrients required by the bacteria [49]. Dumas et al. [22] report that wet manure contains a greater richness and diversity of bacteria than dry litter. Increased moisture is present in the CL, which allows more types of bacteria to persist. However, due to the correlation between moisture content and the levels of other physiochemical parameters known to play a role in microbial diversity, such as carbon and nitrogen availability, determining the predominant factor contributing to bacterial diversity is difficult [22].

The diversity and richness of the bacterial community is also influenced by the pH value; although little data exists regarding this effect, particularly for chicken manure, studies have noted a tendency towards higher community richness and diversity in soils with neutral pH, and lower richness and diversity indexes in more alkaline (pH>8) and acidic (pH<4.5) soils [43]. In our research, CM (pH 8) had higher richness and diversity indexes (*p*≤0.1; trend level) than CL, with a pH value close to neutral pH (pH 6); however, our results are in line with those of Lauber et al. [43] who used samples with similar pH values.

Other differences may result from other factors known to play a crucial role in microbial community structure in chicken waste, like antimicrobial usage, animal diet, nitrogen availability or the presence of heavy metals. Other studies indicate considerable variation in the relative abundance of the bacterial community between chickens, irrespective of the core microbiota colonization [44]. A possible explanation is that shifts in microbial composition are influenced by the initially colonizing microbiota, diet, and immune system of the host [50]. Our present findings indicate an elevated level of calcium in both CL and CM waste. Such high calcium concentrations in poultry diets may result in a reduction in the bioavailability of other minerals, and thus deficiencies in phosphorus, zinc, magnesium and iron. In addition, changes in Ca and P supplementation may alter the composition and activity of the microbial community in the digestive tract of broilers: a high dietary concentration of calcium has been found to increase the pH of crop and ileum contents, but not of the contents of other gastrointestinal tract segments [51]. Furthermore, Ptak et al. [52] report that reductions in Ca and P levels lowered pH and total bacterial count in the caeca; pH value has also been found to directly affect the composition and enzymatic activity of the bacterial community. However, no dietary treatment was found to affect *Bacteroides* counts.

The composition of the microbial community of the crop mucosa was also found to be significantly affected by the Ca and P concentrations in the diet; however, no such effect was observed in digesta samples [44], highlighting the fact that studies on the effects of diet on gut homeostasis should include both digesta and mucosa samples. In addition, Witzig et al. [53] reports that cecal samples possess a different global bacterial community structure to that of the crop, jejunum or ileum, irrespective of the methodological approach, and that this depends on the amount of phosphorus available in the diet. This suggests that the daily intake of P and Ca affects the gastrointestinal microbiome of chickens, as well as their metabolism.

#### 2.2.4. High-throughput quantitative PCR analysis

##### 2.2.4.1. Relative abundance of AMR and MGEs gene classes in HT-qPCR arrays

Among the ARG found in all CM samples, the most abundant were *tet*PB_2 (tetracycline-resistance gene), *tnp*A_2 (MGE), *cmx*A (phenicol-resistance gene), *qac*EΔ1_1 (other; MDR phenotype), *tet*M_3, *tet*X (tetracycline-resistance gene), *qac*EΔ1_2 (other; MDR phenotype), *tet*M_1, and *sul*2_2, *sul*1_1 (sulfonamide-resistance genes); ranked from highest abundance to lowest. In CM, the most abundant group of antimicrobials were tetracycline-resistance genes, with *tet*PB_1, *tet*M_1, and *tet*X being the most prevalent (Figure 5).

**Figure 5.**
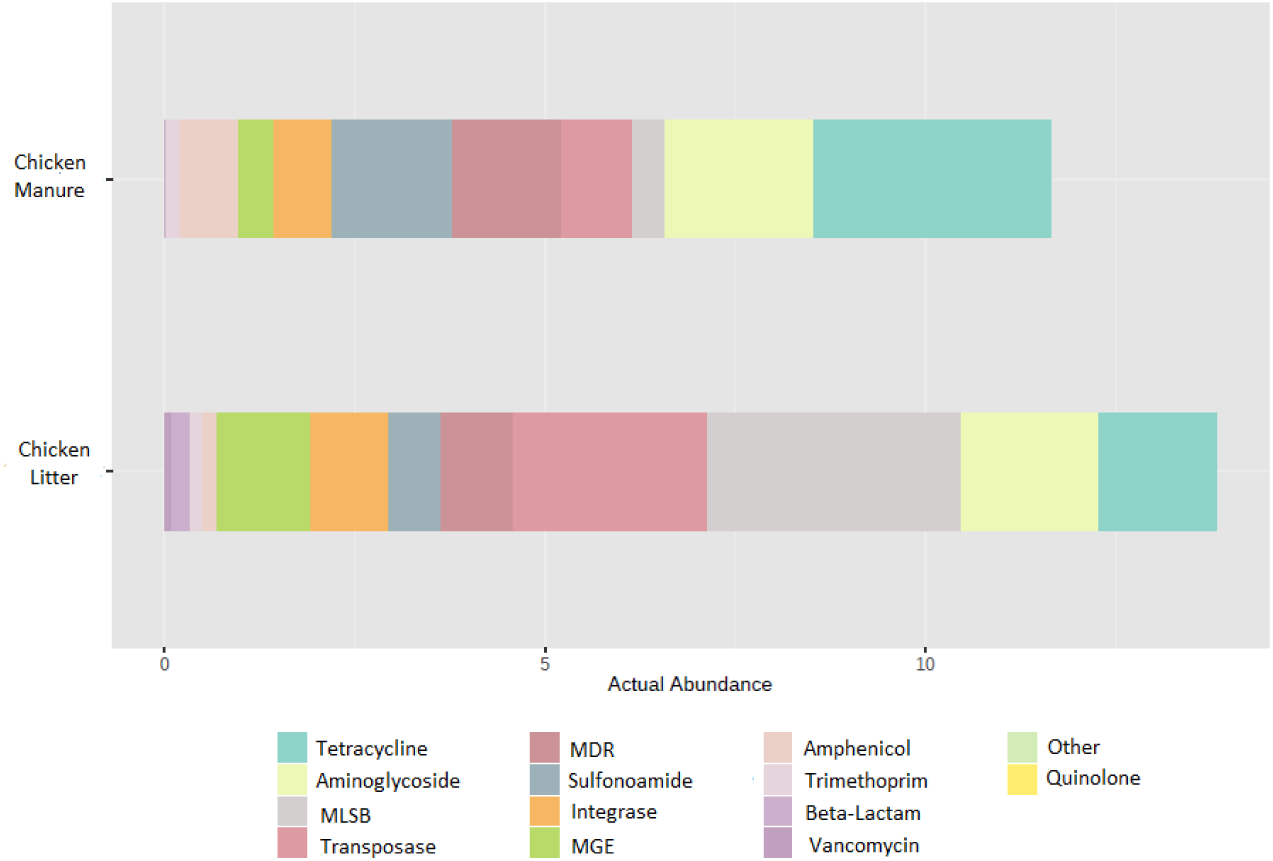
Relative abundance of the detected antibiotic groups, MDR, MGEs, Integrase and Transposase, visualized as gene copy number normalized per 16S rRNA gene copies. The visualization represents the mean value for three replicates.

The most abundant genes in CL were *lnu*A_1 (MLSB-resistance genes), *tnp*A_6, *ISE*fm1 (MGE), *erm*B (MLSB), *tet*M_3 (tetracycline-resistance gene), *int*I1_1 (integrons), *tnp*A_2, *tnp*A_1 (MGE), *tet*M_1 (tetracycline-resistance gene), and *sul*1_1 (sulfonamide-resistance gene). The most prevalent in CL were genes coding resistance to antimicrobials from MLSB (macrolide-lincosamide-streptogramin B) group, with *lnu*A_1, *erm*B, and *erm*X being the most abundant. However, MGE (*tnp*A_6, *ISE*fm1, *tnp*A_2, *tnp*A_1, *pNI105map-F*; ranked from high to low) were even more prevalent. Although CL appears to be the environment with the highest HGT potential (due to high amount of MGE genes), CM can also be classified as an environment of special concern from a microbiological point of view: among the ten most prevalent genes, two are related to an MDR phenotype.

The beta-diversity of the differences in ARGs and MGEs between the two types of waste was visualized by PCoA (Principal Coordinate Analysis) based on Bray-Curtis dissimilarity matrices (Figure 6). Comparisons of the overall distribution patterns of ARGs and MGEs in the samples demonstrated separations between the two groups, i.e. CL and CM, but only at the trend level (*p*<0.1).

**Figure 6.**
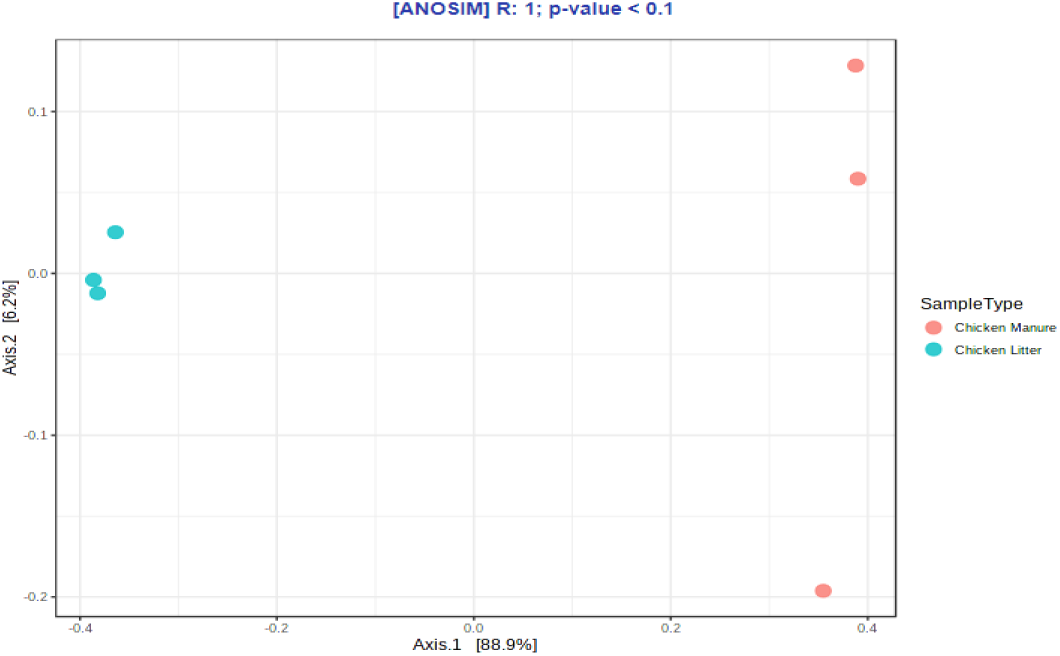
Principal coordinate analysis (PCoA) of ARG and MGE compositions in collected chicken waste samples based on Bray-Curtis dissimilarity matrices (Chicken Manure: CM_1, CM_2,CM_3 [orange dots] and Chicken litter: CL_1, CL_2, CL3 [blue dots]).

The mean distribution of ARG groups found in chicken waste (mean value for CL and CM) directly corresponds to the groups of antibiotics applied to the chickens: polymyxins, penicillin, tetracyclines, trimethoprim and sulfonamides (Figure 7). Also aminoglycosides and antimicrobials belonging to the MLSB group are approved for use in poultry in Poland and by the largest chicken meat producers worldwide (China, Brazil, US) [3]. Tetracycline-resistance genes constitute 19% of all detected ARGs, while aminoglycoside-resistance genes constitute 15%, MLSB-resistance genes 15%, sulfonamide-resistance genes 9%, and other (polymyxins included) 9%. Especially concerning should be the fact that approximately 19% of detected genes belong to MGE, and approximately 7% belong to integrons. This suggests a high potential for antibiotic resistance transfer, which will be analyzed further.

**Figure 7.**
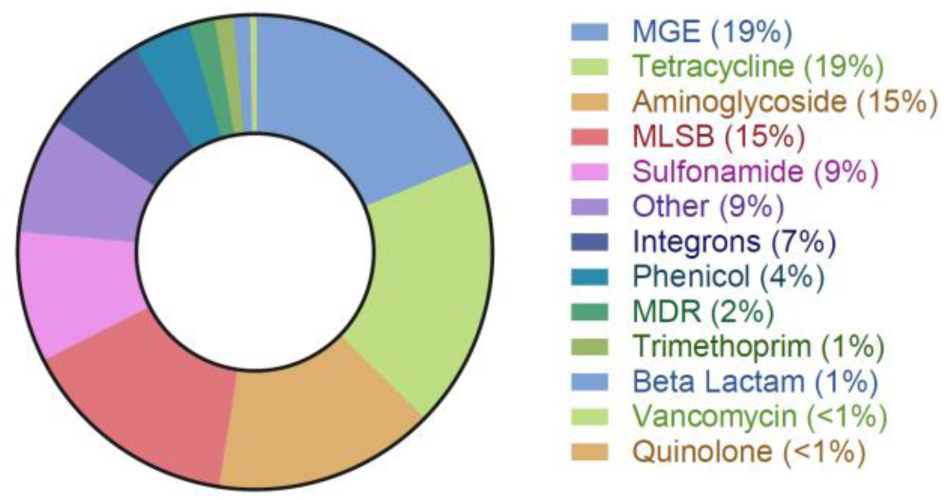
The prevalence of ARG groups, presented as a mean value for all chicken waste samples.

A study of the resistome of raw chicken manure by Peng et al. [54] identified 262 ARGs in chicken manure samples. Similar to our present findings, the main ARGs were aminoglycoside-, tetracycline-, and sulfonamide-resistance genes. In addition, chicken manure was found to be potentially the richest source of ARGs and MGEs of all manures of animal origin [47].

Qian et al. [55] found that all ARGs detected in pig and bovine manure samples were also present in chicken manure. Beta-lactams were found to be the most abundant ARGs in bovine manure (14.4–68.2%), while only residual amounts were present in the two other manure types. This is in line with our present results, where the level of beta-lactam abundance was low in CL samples (0.07), and even lower in CM (0.004). As also reported by Qian et al. [55], MGEs comprising *Int*I-1, *Tp*614, *IS*613, and *tnp*A were detected in all our tested manure samples.

In other studies conducted by Xu et al. [56], chicken manure was fond to demonstrate higher ARG abundance than other examined sheep and cattle manure. In the chicken manure samples, similar to ours, the most abundant ARG was found to be *tet*X, probably due to the broad range of potential hosts. In addition, the *sul*1 and *sul*2 genes were pre-existing and dominant in those samples, and importantly, their numbers considerably decreased during the composting process [56].

Out of the 384 tested ARGs and MGEs, 162 ARGs were identified in CM samples and 237 in CL (Figure S1, Figure 8). Core resistome analysis revealed that CM and CL samples share 141 genes in common, and that CL has 96 unique genes and CM has 21. The CL samples have both a higher actual abundance of ARG (Figure 5) and the highest diversity of detected genes. Among the most prevalent genes in CL and CM, only *tet*PB_2 exists exclusively in CM; the remainder are common for both groups of chicken waste, but exhibit lower frequencies (*tnp*A_2, *cmx*A, *qacE*Δ1_1, *tet*M_3, *tet*X, *qac*EΔ1_2, *tet*M_1, *sul*1_1, *sul*1_2, *lnu*A_1, *tnp*A_6, *ISE*fm1, *erm*B, *int*I1_1, *tnp*A_2, *tnp*A_1, *tet*M_1).

**Figure 8.**
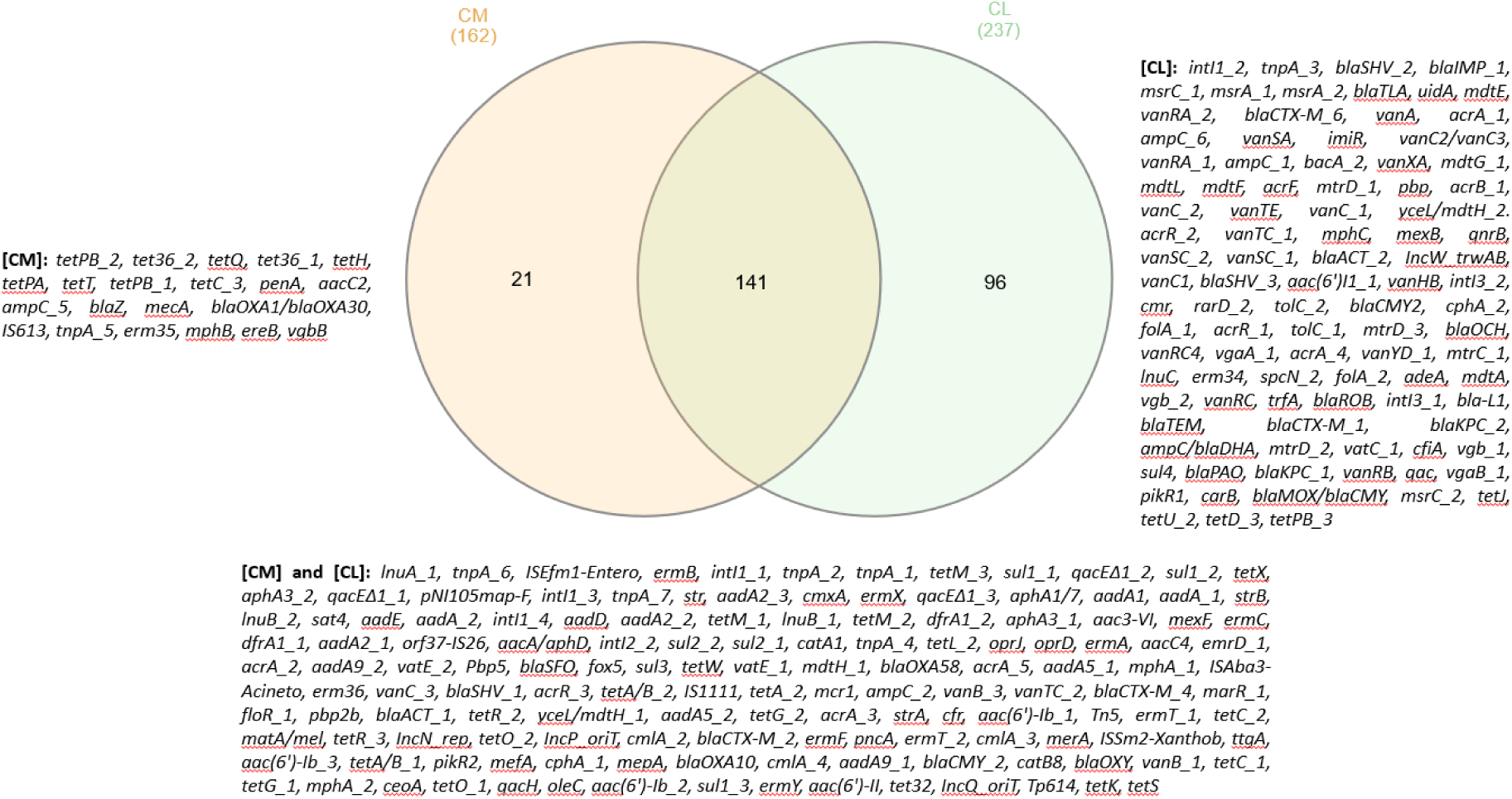
Comparison between CL and CM resistomes.

Recent studies comparing human, pig, and chicken gut resistomes that detected a total of 166 ARGs identified the highest ARG levels in the chicken samples (156 ARGs). In addition, although the chicken samples demonstrated the greatest overall abundance, their ARGs had the lowest diversity [57]. This high abundance of ARGs in chicken might be connected with high breeding density and their short, intense growth period [58]. However, several ARGs and MGEs were also found to be present on farms with no history of antibiotic usage; similarly to our present samples, the most prevalent ARGs in samples from the antibiotic-free farms were the aminoglycoside-resistance genes *aad*A, *aad*A1, *aad*A2, and *str*B, sulfonamide-resistance gene *sul*2, and tetracycline-resistance genes *tet*M, *tet*K and *tet*X [59].

##### 2.2.4.2. Correlation analysis of ARGs, MGEs and microbial communities

###### Correlation between ARGs and MGEs

An ARG and MGE co-occurrence network based on Spearman’s correlations was constructed to analyze the potential risk of spread of antimicrobial resistance into the environment via HGT (Figure 9). This network was based on 1068 significant correlations (46 negative and 1022 positive) and contained 218 nodes (25 MGEs and 193 ARGs) and 1069 edges. As the size of the nodes is related to the degree of its interactions, our network indicates that MGEs play a crucial role. Among all the detected MGEs, *Inc*W_*trw*AB and *Inc*Q_*ori*T (plasmid incompatibility) and *trf*A (transposase) were found to have largest number of significant interactions with ARGs. In addition, two MGEs were found to have a negative interaction with tetracycline-resistance genes: *tnpA*_3 (transposase) and *pNI105map-F* (plasmid replication) – the former with *tet*R_3, *tet*G_2, and *tet*R_2 and the latter with *tet*M_2, *tet*36_2, and *tet*PA. The only gene identified in the tested samples encoding quinolone-resistance (*qnr*B) only demonstrated positive interactions with nine MGEs (e.g., *int*I3_1, *Inc*N_rep). *Sul*1_2 showed negative correlation with the transposase gene *tnp*A_3, and *sul*1_3 with *tnp*A_7.

**Figure 9.**
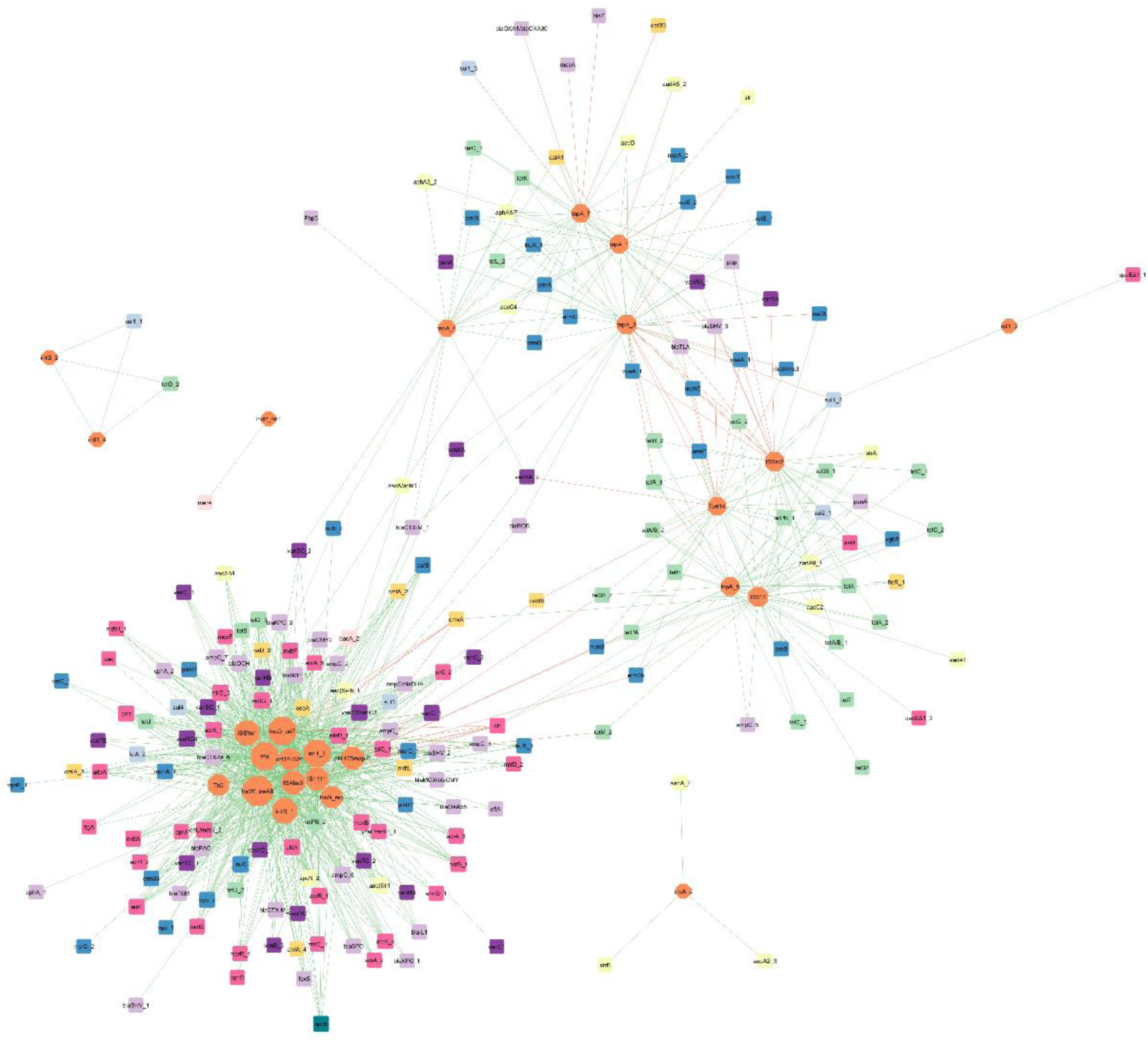
ARG and MGE interaction network analysis presented in ‘organic layout’. A connection shows a strong and significant correlation based on Spearman’s rank analysis (|r|≥0.9, *p*<0.01). The red and green edges indicated the indexes of positive and negative correlations between ARGs and MGEs, respectively. The size of the nodes showed the degree of the interactions.

Among the beta-lactam resistance genes, *tnp*A_7 (transposase) gene is negatively correlated with three genes: *bla*Z, *bla*OXA1/*bla*OXA30, and *mec*A (methicillin-resistance). Two different genes, viz. *Tp*614 (transposase) and *ISSm2* (insertional sequence), demonstrate negative interactions with *bla*TLA, *bla*SHV_3 genes, with the latter having an additional relationship with the *pbp* gene. The only negative interaction between the aminoglycoside-resistance genes and MGEs is that between *aad*A5_2 and *tnp*A_1 (transposase). Furthermore, in this network, a negative correlation can be seen between *tnp*A_3 and two other MGEs (*ISSm2*-insertional sequence and *Tp*614-transposase).

Several negative correlations were found between vancomycin-resistance genes and MGEs such as these encoding the insertional sequence *ISSm2* and *Tp*614; the former is negatively correlated with two genes *van*SA and *van*RA_1 and the latter with *van*RA_2. Out of the several positive interactions between genes encoding the MLSB phenotype, the *pNI105map-F* gene is negatively correlated with *mph*B (macrolide) and *erm*35 (antibiotic target alteration). Similarly, the *mph*C gene has a negative correlation with *ISS*m2 and *Tp*614. Additionally *Tp*614 is negatively correlated with *msr*A_1 (ABC-F ATP-binding cassette ribosomal protection protein). The gene *erm*Y (a plasmid-mediated methyltransferase) is correlated negatively with two variants of transposase *tnp*A_1 and *tnp*A_3; the latter also interacts negatively with *mat*A/mel genes. For the *cfr* gene (ribosomal RNA methyltransferase), six out of eight significant interactions with MGEs were found to be negative (e.g., *Tn*5, *IS*1111, *trf*A).

Such network analysis can play an important role in understanding the role of MGEs in the transfer of antimicrobial resistance to the environment and can provide new information about possible spreading mechanisms and patterns. One of the most prevalent groups of genes in manure and soil is the MLSB resistance genes, such as erythromycin ribosome methylation (*erm*) [54,60]. Zhang et al. [11] studied the succession of soil antibiotic resistance following the application of three types of animal manure; the results indicate a significant correlation between three erm genes (*erm*A, *erm*B, *erm*C), with transposons indicating their potential transfer [11]. In our studies, these *erm* genes were found to interact significantly with *tnp*A (transposase) gene subtypes.

Another group of antimicrobials widely used in veterinary scenarios are beta-lactams. They are often applied to treat diseases, such as mastitis, and for prophylaxis in animals. Extended-spectrum beta-lactamase (ESBL) producers can be isolated from different environments (soil, water, animal manure) [61]. Evidence exists supporting the possible spread of ESBL genes by HGT; for example, *bla*TEM and *bla*CTX-M-1 genes detected in pig manure samples have been associated with IncN plasmids [62,63]. In our network analyses found *bla*CTX-M-1 to interact with two types of genes (I) introns (*intI*1_2, *int*I3_2) and (II) transposases (*tnp*A_3, *tnp*A_6), and only *bla*TEM interacted with the *Inc*N_*rep* gene.

Two groups of ARGs known to be prevalent in manures are tetracycline and sulfonamides-resistance genes. Due to their low absorbance in the gastric tract, most of the administered drug is exerted as an active compound with feces and urine into the environment [64]. Hence, these resistance genes are present in high amounts in fresh manure samples and manure-treated soils [65–67]. Various types of *tet* genes present in animal manure are inserted in transposons [68,69]. Most detected *tet* genes in our network are clustered and connected with *Tp*614, *IS*614 and *ISS*m2 (insertional sequences) and *tnp*A_3 (transposase) genes.

Sulfonamide-resistance genes (*sul*1, *sul*2, *sul*3) are widely present in many natural environments like soil and animal manure. They can be located on MGEs and transfer between non-related bacteria by mobilized broad host plasmids [70]. *Sul*1 and *sul*2 are frequently associated with 1 class integrons (intI) and hence can be easily transferred [70,71]. Our network confirmed a strong and significant association between *sul* and int genes. *Sul*1_1 genes interact only with *int*I2_2 and *int*I1_4 subtypes. *Sul*1_2 and *qac*EΔ1_1 interact with the *int*I1_3 gene. Most importantly, *sul*2_1 and *sul*1_2 genes are also connected with *Tp*614, *IS*614 and *ISS*m2 (insertional sequences).

The only gene connected to the *Inc*P_*ori*T gene (plasmid) is the mercury reductase gene (*mer*A). The presence of this gene was confirmed to be strongly connected with presence of IncP-α plasmid in High Arctic snow, freshwater and sea-ice brine [72]. Plasmids from this incompatibility group can spread resistance genes among even non-related phylogenetic groups of Gram-negative bacteria [73].

###### Correlation between detected genes and microbial taxa at phylum level

The second network, showing the interactions between nine taxa at phylum level and eight gene classes (seven ARG classes and MGEs encoding genes) in chicken waste, is based on 79 strong and significant correlations (41 positive and 38 negative). The network was formed of 17 nodes (eight antibiotic groups and MGEs and nine microbial taxa at the phylum level). According to the network, organized according to Spearman’s rank analysis (|r|≥0.7, *p*<0.05), various ARGs appear to be significantly associated with different bacterial phyla.

The potential host microorganisms that were positively correlated with ARGs mainly belonged to *Actinobacteria, Proteobacteria* and *Firmicutes. Bacteroidetes* had significant positive interaction only with sulfonamide- and tetracycline-resistance genes. When it comes to antimicrobial groups, quinolone-resistance genes had a significant positive correlation with only two phyla: *Actinobacteria* and *Firmicutes*. Amphenicol-resistance genes had a negative correlation with almost all taxa, except *Actinobacteria*. Sulfonamide-resistance genes had a negative correlation with *Chloroflexi, Bacteroidetes* and *Tenericutes*. MGEs showed positive correlations with *Proteobacteria, Actinobacteria* and *Firmicutes* (three phyla characterized as dominant in examined samples) (Figure 10). Similar effects were presented by Zhou et al. [74], which identified the same potential hosts of ARGs as in our analyses, belonging to the four most dominant phyla identified in the samples (*Firmicutes, Proteobacteria, Bacteroidetes*, and *Actinobacteria*).

**Figure 10.**
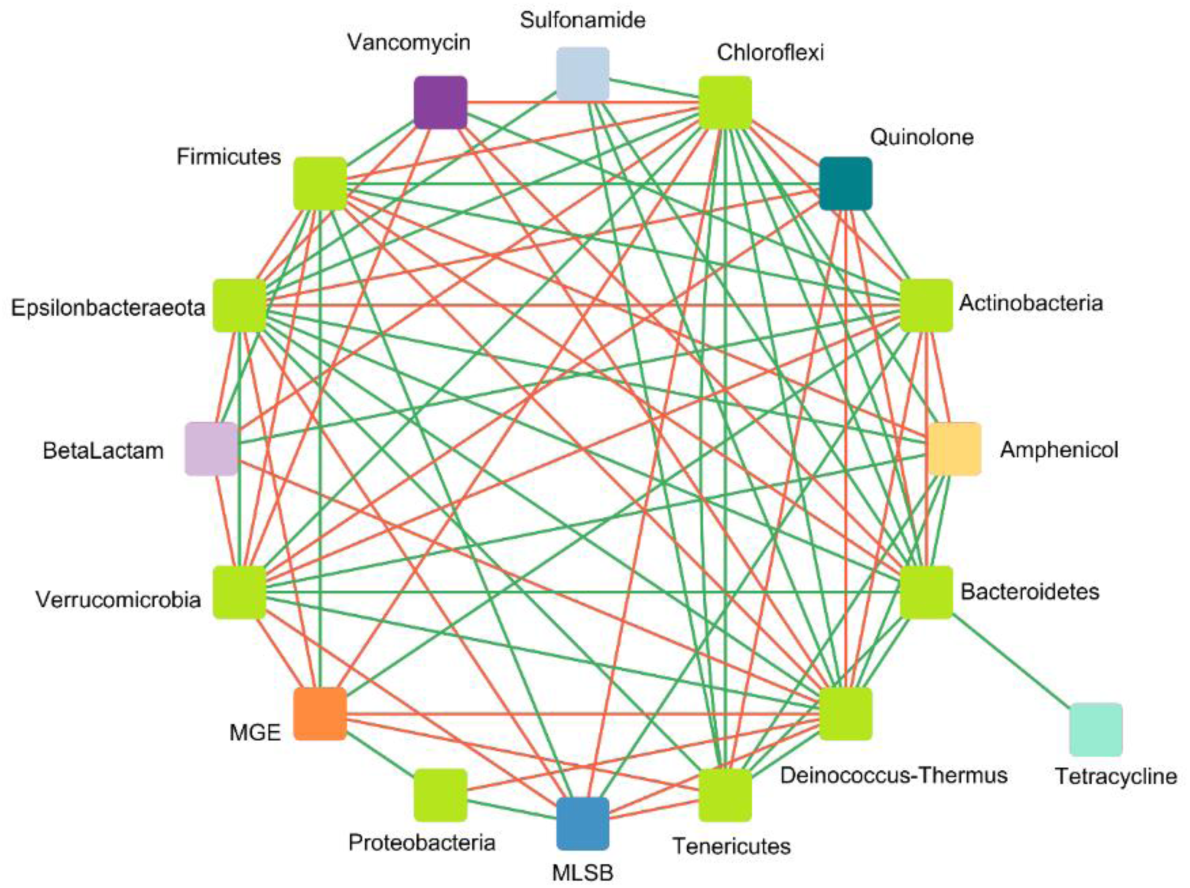
Microbial taxon and gene class interaction network analysis as a ‘circular layout’ based on the co-occurrence of ARG groups and microbial taxa at phylum level. A connection shows a significant correlation based on Spearman’s rank analysis (|r|≥0.7, *p*<0.05). The green and red edges indicate positive and negative correlations between ARGs and taxa, respectively.

In other studies conducted on composting chicken manure, most of the positive interactions with ARGs were noted for *Proteobacteria*, e.g., *Pseudomonas, Halomonas* and *Stenotrophomonas*. This group demonstrated a significant correlation with the *tet*O, *tet*C and *sul*1 genes [75]. In our samples, the ARGs coding resistance to tetracyclines only displayed a significant and positive correlation with *Bacteroidetes*. In animal manure, more than a half of the significant correlation with ARGs and MGEs was attributed to *Proteobacteria* and *Bacteroidetes*; these were also the dominant phyla in our analyzed samples [76].

## 3. Materials and Methods

### 3.1. Manure sample

Chicken waste samples were collected from two different sources of meat production at a large, commercial, meat-producing farm in Poland (CM - chicken manure from laying hens and CL - chicken litter from broilers). The farm owner agreed to manure sampling. The samples were transported immediately after collection to the laboratory, stored at 4 °C, and processed within 24 hours. The samples were subjected to physicochemical analyses at the Wessling Poland accredited analytical laboratory; these analyses comprised pH value, biogenic element, and heavy metal content: Hg content according to procedure DIN ISO 16772, other heavy metals (ICP-OES/ICP-MS) – DIN EN ISO 11885/DIN EN ISO 17294-2, total nitrogen – VDLUFA, Bd. I, Kap. A 2.2.1, ammoniacal nitrogen – DIN 38406 E5-1 mod., dry organic matter - DIN EN 12879. Only Hg content and dry organic matter content determination were performed according to the accredited norms: other components were analyzed according to procedures described in the norms, but adjusted to our needs.

### 3.2. Bacterial strain isolation and characterization

Prior to isolating the microorganisms of interest, the obtained waste was enriched with Brain Heart Infusion Broth (for *Enterococcus* spp.) or Luria Bertani Broth (Tab. S2) by adding 1g of chicken waste to 9 ml of liquid medium. The cultures were incubated at 25 °C and 37 °C for 24 h and 48 h, diluted (10^−1^, 10^−2^, 10^−3^), and then 100 µl of each dilution was plated on an appropriate selective medium with antibiotic supplementation according to Table S2. The media and antibiotics were chosen according to the WHO priority pathogens list [21]. Each isolation was performed as three biological replicates followed by three technical ones. All selective media were purchased from Biomaxima, Poland, except for Brillance VRE Agar, which was obtained from OXOID. Antibiotics and inositol were purchased from Sigma-Aldrich (MERCK).

The plates were then incubated according to the recommendations of the medium manufacturer and again at 25 °C to select for bacteria originating from the natural environment. Following this, approximately 24-48 colonies resembling the morphology of the intended bacteria were isolated from each medium type, starting from the plate with the highest dilution. Pure cultures were obtained by three consecutive streaking. The resulting cultures were considered pure and stored for further analysis in PBS/glycerol stocks (20% v/v).

The Kirby-Bauer test for determining antibiotic susceptibility was applied to assess the phenotypic diversity of isolated strains and eliminate any potentially repetitive strains [77]. Antimicrobial discs (OXOID) were chosen according to the selective medium used for isolation (Table S3). After incubation for 18 hours at 37 °C, the inhibition zones were measured. Strains with different antimicrobial susceptibility profiles (inhibition zone varied by approximately +/- 2 mm in diameter) around at least one antibiotic disk were considered nonrepetitive and chosen for further research.

The strains isolated from the chicken waste were classified to species, or genus, level using a matrix-assisted laser desorption/ionization system equipped with a time-of-flight mass spectrometer (MALDI-TOF MS/MS) [78]. The analysis was conducted in an accredited external laboratory, ALAB Laboratoria Sp. z o.o., Poland, according to a standard diagnostic procedure. Identification was performed by aligning the peaks to the best matching reference data; the resulting log score was classified as follows: ≥2.3, highly probable species; between 2.0 and 2.3, certain genus and probable species; between 1.7 and 2.0, probable genus; and <1.7, non-reliable identification.

Where applicable, antibiotic susceptibility testing was performed using the VITEK® 2 Compact System (BioMerieux, France) [79]. Several antimicrobial susceptibility test cards (AST Cards) were used depending on the identified bacteria species (AST-P643, AST-ST03, AST-P644, AST-N331, AST-N332). Each susceptibility test provides an accurate susceptibility phenotype profile for almost all tested bacterial strains. Based on The European Committee on Antimicrobial Susceptibility Testing (EUCAST) recommendations (EUCAST, Breakpoint tables for interpretation of MICs and zone diameters, 2020), bacterial strains were assigned as resistant (R), sensitive (S), and intermediate (I) to applied antibiotics. Standard disk diffusion method was used for isolates where automatic testing was not provided due to the lack of the EUCAST recommendations for this particular bacterial species/genus. Strains were considered resistant only when no inhibition zone appeared around the antibiotic discs. In all other cases, strains were considered sensitive.

### 3.3. DNA extraction and metagenomic sequencing

DNA extraction was performed from 0.5 g of each type of waste with the FastDNA(tm) SPIN Kit for Feces and the FastPrep Homogenizer (MP Biomedicals, USA) according to the manufacturer’s instructions. The DNA concentration was determined by Qubit fluorometer and dsDNA BR Assay Kit (Thermo Fisher Scientific, USA), and purity was determined by measuring the A260/A280 absorbance ratio with a NanoDrop (Thermo Fisher, USA). Only samples with concentrations higher than 200 µg/ml and A260/A280 ratio ranging from 1.8 to 2.0 were chosen for analysis. DNA samples were stored at -20 °C for further use. The DNA samples were isolated in triplicate.

The microbial biodiversity of waste samples was determined by sequencing targeting the variable V3-V4 regions of the bacterial 16S rRNA gene using the Illumina platform. Libraries were prepared with the Nextera® XT Library Preparation Kit (Illumina) according to the manufacturer’s recommendations. The obtained datasets were analyzed using Qiime 2 pipeline with the DADA2 option (sequence quality control) and SILVA ribosomal RNA amplicon database (taxonomy assigned) [80]. Targeted metagenome sequencing was performed on the Illumina MiSeq platform at the Institute of Biochemistry and Bio-physics, Polish Academy of Sciences (IBB, Poland). The obtained data were analyzed and visualized using the MicrobiomeAnalyst web tool [81].

### 3.4. qPCR analysis of the ARG profile

In all chicken waste samples, the presence of ARG, integrons and MGE was analyzed by a high-throughput SmartChip qPCR system (Resistomap, Finland). Their amounts were calculated as a relative number of copies (gene copy numbers per 16S rRNA gene copy numbers). The qPCR conditions and initial data processing were performed as described previously [70]. All qPCR reactions were done in three repetitions, and a threshold cycle (Ct) of 27 was used as the detection limit. Only samples with two or three replicates and the parameters mentioned earlier (Ct≤27, efficiency 1.8-2.2) were regarded as positive. Genes and primers are listed in Supplementary materials file 1. Further, gene names and gene groups were used across the manuscript according to those listed and categorized in this table.

The relative copy number was calculated as follows: Relative gene copy number = 10^(27-Ct)/(10/3)^ [82]. The ARG copy numbers were calculated by normalizing the relative copy numbers per 16S rRNA gene copy numbers.

Spearman’s correlation coefficient (together with all data visualization) was calculated with GraphPad Prism 9 to identify the relationships between ARGs, MGEs, and microbial taxa. The analyzed relationships were used to construct networks using Cytoscape. The correlations were considered strong and significant when the absolute value of Spearman’s rank |r|≥0.9 and *p*<0.01 (unless stated otherwise).

## 4. Conclusions

Our findings indicate that the examined chicken waste can be a sufficient source of nutrients essential for plant growth like nitrogen and phosphorus, especially considering that the amounts of heavy metals in the chicken manures were within acceptable levels and would not likely pose an environmental threat when directly applied to the soil. Thus taking into consideration only physicochemical parameters of chicken wastes, these type of manure can be apply on arable fields as a natural fertilizer with high level nutrition potential. The microbial community composition analyses found the dominant bacterial phyla in chicken waste to be *Bacteroidetes, Firmicutes*, and *Proteobacteria*. Our data also found the diversity and richness of microbial communities to be associated with the physical and chemical conditions of the collected samples.

During the study, we identified an alarming diversity and abundance of ARGs and MGEs, and our data indicates positive correlations between them suggesting a considerable risk of ARG spread into the environment and their transfer between bacterial species. The most prevalent antibiotic resistance genes in both types of studied chicken waste, viz. chicken litter (CL) and chicken manure (CM), were aminoglycoside-, sulfonamide- and tetracycline-resistance genes. In CL, genes related to the MLSB phenotype predominate. Interestingly, CL and CM were found to differ when comparing the overall distribution patterns of antibiotic-resistance genes and mobile genetic elements.

Moreover, we isolated antibiotic resistant bacteria strains, some of them were potentially pathogenic, such as *Staphylococcus lentus*, while others were considered opportunistic human and animal pathogens, such as *Myroides odoratus* or *M. odoratimimus*, or phytopathogens, like *Flavobacterium johnsoniae*. Even more concerning is that many of them were resistant to at least one antibiotic class, and more than 50% of isolated strains were classified as MDR, due to the resistance against more than three antibiotic classes. Hence, it can be concluded that the opportunistic pathogens with MDR phenotypes present in chicken manure and litter pose a real risk of reaching the ground and groundwater and contaminating the surrounding environment, which poses a potential risk to public health. Therefore, taking into account all the results obtained, it is fair to qualify the studied chicken waste as highly dangerous material.

## Author Contributions

A.B.# – Investigation, Formal Analysis, Writing - Original Draft, Visualization; M.Z.# – Investigation, Methodology, Formal Analysis, Writing - Original Draft, Visualization; A.G. - Methodology, Writing - Review & Editing, M.P. – Conceptualization, Writing - Review & Editing, Supervision, Funding acquisition, Project administration; #authors contributed equally

## Data Availability Statement

Datasets used during the study have been shared as Supporting materials.

## References

1. FAO FAOSTAT Available online: https://www.fao.org/faostat/en/#home (accessed on 23 March 2022).

2. Muhammad, J.; Khan, S.; Su, J.Q.; Hesham, A.E.L.; Ditta, A.; Nawab, J.; Ali, A. Antibiotics in Poultry Manure and Their Associated Health Issues: A Systematic Review. Journal of Soils and Sediments: protection, risk assessment and remediation 2020, 20, 486–497, doi:10.1007/s11368-019-02360-0.

3. Roth, N.; Käsbohrer, A.; Mayrhofer, S.; Zitz, U.; Hofacre, C.; Domig, K.J. The Application of Antibiotics in Broiler Production and the Resulting Antibiotic Resistance in Escherichia Coli: A Global Overview. Poult. Sci. 2019, 98, 1791–1804, doi:10.3382/ps/pey539.

4. Berendsen, B.J.A.; Lahr, J.; Nibbeling, C.; Jansen, L.J.M.; Bongers, I.E.A.; Wipfler, E.L.; van de Schans, M.G.M. The Persistence of a Broad Range of Antibiotics during Calve, Pig and Broiler Manure Storage. Chemosphere 2018, 204, 267–276, doi:10.1016/j.chemosphere.2018.04.042.

5. Zhang, Y.; Geissen, S.-U.; Gal, C. Carbamazepine and Diclofenac: Removal in Wastewater Treatment Plants and Occurrence in Water Bodies. Chemosphere 2008, 73, 1151–1161, doi:10.1016/j.chemosphere.2008.07.086.

6. Poole, T.; Sheffield, C. Use and Misuse of Antimicrobial Drugs in Poultry and Livestock: Mechanisms of Antimicrobial Resistance. Pakistan Veterinary Journal 2013, 33, 266–271.

7. Archawakulathep, A.; Kim, C.T.T.; Meunsene, D.; Handijatno, D.; Hassim, H.A.; Rovira, H.R.G.; Myint, K.S.; Baldrias, L.R.; Sothy, M.; Aung, M.; et al. Perspectives on Antimicrobial Resistance in Livestock and Livestock Products in ASEAN Countries. The Thai Journal of Veterinary Medicine 2014, 44, 5–13.

8. Rothrock, M.J.; Hiett, K.L.; Guard, J.Y.; Jackson, C.R. Antibiotic Resistance Patterns of Major Zoonotic Pathogens from All-Natural, Antibiotic-Free, Pasture-Raised Broiler Flocks in the Southeastern United States. J Environ Qual 2016, 45, 593–603, doi:10.2134/jeq2015.07.0366.

9. Franklin, A.M.; Aga, D.S.; Cytryn, E.; Durso, L.M.; McLain, J.E.; Pruden, A.; Roberts, M.C.; Rothrock, M.J.; Snow, D.D.; Watson, J.E.; et al. Antibiotics in Agroecosystems: Introduction to the Special Section. J Environ Qual 2016, 45, 377–393, doi:10.2134/jeq2016.01.0023.

10. Zalewska, M.; Błażejewska, A.; Czapko, A.; Popowska, M. Antibiotics and Antibiotic Resistance Genes in Animal Manure – Consequences of Its Application in Agriculture. Frontiers in Microbiology 2021, 12.

11. Zhang, Y.-J.; Hu, H.-W.; Gou, M.; Wang, J.-T.; Chen, D.; He, J.-Z. Temporal Succession of Soil Antibiotic Resistance Genes Following Application of Swine, Cattle and Poultry Manures Spiked with or without Antibiotics. Environ Pollut 2017, 231, 1621–1632, doi:10.1016/j.envpol.2017.09.074.

12. Drózdz, D.; Wystalska, K.; Malińska, K.; Grosser, A.; Grobelak, A.; Kacprzak, M. Management of Poultry Manure in Poland – Current State and Future Perspectives. Journal of Environmental Management 2020, 264, 110327, doi:10.1016/j.jenvman.2020.110327.

13. Wang, L.; Lilburn, M.; Yu, Z. Intestinal Microbiota of Broiler Chickens As Affected by Litter Management Regimens. Frontiers in Microbiology 2016, 7.

14. Lovanh, N.; Cook, K.L.; Rothrock, M.J.; Miles, D.M.; Sistani, K. Spatial Shifts in Microbial Population Structure within Poultry Litter Associated with Physicochemical Properties. Poult Sci 2007, 86, 1840–1849, doi:10.1093/ps/86.9.1840.

15. Collins, E.; Barker, J.C.; Carr, L.E.; Brodie, H.L.; Martin, J.H. Poultry Waste Management Handbook (NRAES 132 - FRONT MATTER ONLY); Natural Resource, Agriculture, and Engineering Service (NRAES), 1999;

16. Manyi-Loh, C.; Mamphweli, S.; Meyer, E.; Okoh, A. Antibiotic Use in Agriculture and Its Consequential Resistance in Environmental Sources: Potential Public Health Implications. Molecules 2018, 23, doi:10.3390/molecules23040795.

17. Laconi, A.; Mughini-Gras, L.; Tolosi, R.; Grilli, G.; Trocino, A.; Carraro, L.; Di Cesare, F.; Cagnardi, P.; Piccirillo, A. Microbial Community Composition and Antimicrobial Resistance in Agricultural Soils Fertilized with Livestock Manure from Conventional Farming in Northern Italy. Sci Total Environ 2021, 760, 143404, doi:10.1016/j.scitotenv.2020.143404.

18. Lima, T.; Domingues, S.; Da Silva, G.J. Manure as a Potential Hotspot for Antibiotic Resistance Dissemination by Horizontal Gene Transfer Events. Veterinary Sciences 2020, 7, 110, doi:10.3390/vetsci7030110.

19. Hu, Y.; Yang, X.; Li, J.; Lv, N.; Liu, F.; Wu, J.; Lin, I.Y.C.; Wu, N.; Weimer, B.C.; Gao, G.F.; et al. The Bacterial Mobile Resistome Transfer Network Connecting the Animal and Human Microbiomes. Appl Environ Microbiol 2016, 82, 6672–6681, doi:10.1128/AEM.01802-16.

20. Chen, Z.; Jiang, X. Microbiological Safety of Chicken Litter or Chicken Litter-Based Organic Fertilizers: A Review. Agriculture 2014, 4, 1–29, doi:10.3390/agriculture4010001.

21. Tacconelli, E.; Carrara, E.; Savoldi, A.; Harbarth, S.; Mendelson, M.; Monnet, D.L.; Pulcini, C.; Kahlmeter, G.; Kluytmans, J.; Carmeli, Y.; et al. Discovery, Research, and Development of New Antibiotics: The WHO Priority List of Antibiotic-Resistant Bacteria and Tuberculosis. The Lancet Infectious Diseases 2018, 18, 318–327, doi:10.1016/S1473-3099(17)30753-3.

22. Dumas, M.D.; Polson, S.W.; Ritter, D.; Ravel, J.; Jr, J.G.; Morgan, R.; Wommack, K.E. Impacts of Poultry House Environment on Poultry Litter Bacterial Community Composition. PLOS ONE 2011, 6, e24785, doi:10.1371/journal.pone.0024785.

23. Dikinya, O.; Mufwanzala, N. Chicken Manure-Enhanced Soil Fertility and Productivity: Effects of Application Rates. 2010, doi:10.5897/JSSEM.9000019.

24. Regulation (EC) No 1069/2009 of the European Parliament and of the Council of 21 October 2009 Laying down Health Rules as Regards Animal by-Products and Derived Products Not Intended for Human Consumption and Repealing Regulation (EC) No 1774/2002 (Animal by-Products Regulation); 2009; Vol. 300;.

25. Rogeri, D.; Ernani, P.; Mantovani, A.; Lourenço, K. Composition of Poultry Litter in Southern Brazil. Revista Brasileira de Ciência do Solo 2016, 40, doi:10.1590/18069657rbcs20140697.

26. Sarbjit Singh, G.S.; Shamsuddin, M.R.; Aqsha Lim, S. Characterization of Chicken Manure from Manjung Region. IOP Conference Series: Materials Science and Engineering 2018, 458, 012084, doi:10.1088/1757-899X/458/1/012084.

27. Li, G.; Li, H.; Leffelaar, P.A.; Shen, J.; Zhang, F. Characterization of Phosphorus in Animal Manures Collected from Three (Dairy, Swine, and Broiler) Farms in China. PLOS ONE 2014, 9, e102698, doi:10.1371/journal.pone.0102698.

28. Reyer, H.; Oster, M.; Ponsuksili, S.; Trakooljul, N.; Omotoso, A.O.; Iqbal, M.A.; Muráni, E.; Sommerfeld, V.; Rodehutscord, M.; Wimmers, K. Transcriptional Responses in Jejunum of Two Layer Chicken Strains Following Variations in Dietary Calcium and Phosphorus Levels. BMC Genomics 2021, 22, 485, doi:10.1186/s12864-021-07814-9.

29. Proszkowiec-Weglarz, M.; Angel, R. Calcium and Phosphorus Metabolism in Broilers: Effect of Homeostatic Mechanism on Calcium and Phosphorus Digestibility1 1Presented as a Part of the Informal Nutrition Symposium “Metabolic Responses to Nutrition and Modifiers” at the Poultry Science Association’s Annual Meeting in Athens, Georgia, July 9, 2012. Journal of Applied Poultry Research 2013, 22, 609–627, doi:10.3382/japr.2012-00743.

30. Bedford, M.; Rousseau, X.; Bedford, M.; Rousseau, X. Recent Findings Regarding Calcium and Phytase in Poultry Nutrition. Anim. Prod. Sci. 2017, 57, 2311–2316, doi:10.1071/AN17349.

31. Shafey, T.M. Calcium Tolerance of Growing Chickens: Effect of Ratio of Dietary Calcium to Available Phosphorus. World’s Poultry Science Journal 1993, 49, 5–18, doi:10.1079/WPS19930002.

32. Meyer, A.; Dang, H.; Roland, W. Myroides Spp. Cellulitis and Bacteremia: A Case Report. IDCases 2019, 18, e00638, doi:10.1016/j.idcr.2019.e00638.

33. Ktari, S.; Mnif, B.; Koubaa, M.; Mahjoubi, F.; Ben Jemaa, M.; Mhiri, M.N.; Hammami, A. Nosocomial Outbreak of Myroides Odoratimimus Urinary Tract Infection in a Tunisian Hospital. J Hosp Infect 2012, 80, 77–81, doi:10.1016/j.jhin.2011.09.010.

34. Ewers, C.; Antão, E.-M.; Diehl, I.; Philipp, H.-C.; Wieler, L.H. Intestine and Environment of the Chicken as Reservoirs for Extraintestinal Pathogenic Escherichia Coli Strains with Zoonotic Potential. Appl Environ Microbiol 2009, 75, 184–192, doi:10.1128/AEM.01324-08.

35. Marano, R.B.M.; Fernandes, T.; Manaia, C.M.; Nunes, O.; Morrison, D.; Berendonk, T.U.; Kreuzinger, N.; Telson, T.; Corno, G.; Fatta-Kassinos, D.; et al. A Global Multinational Survey of Cefotaxime-Resistant Coliforms in Urban Wastewater Treatment Plants. Environment International 2020, 144, 106035, doi:10.1016/j.envint.2020.106035.

36. Moffo, F.; Mouiche, M.M.M.; Djomgang, H.K.; Tombe, P.; Wade, A.; Kochivi, F.L.; Dongmo, J.B.; Mbah, C.K.; Mapiefou, N.P.; Ngogang, M.P.; et al. Poultry Litter Contamination by Escherichia Coli Resistant to Critically Important Antimicrobials for Human and Animal Use and Risk for Public Health in Cameroon. Antibiotics 2021, 10, 402, doi:10.3390/antibiotics10040402.

37. Agostinho, J.M.A.; Cardozo, M.V.; Borzi, M.M.; Marin, J.M. Antibiotic Resistance and Virulence Factors among Escherichia Coli Isolates from Avian Organic Fertilizer. Cienc. Rural 2020, 50, e20180849, doi:10.1590/0103-8478cr20180849.

38. Moraru, R.; Pourcher, A.-M.; Jadas-Hecart, A.; Kempf, I.; Ziebal, C.; Kervarrec, M.; Comunal, P.-Y.; Mares, M.; Dabert, P. Changes in Concentrations of Fluoroquinolones and of Ciprofloxacin-Resistant Enterobacteriaceae in Chicken Feces and Manure Stored in a Heap. J Environ Qual 2012, 41, 754–763, doi:10.2134/jeq2011.0313.

39. Atidégla, S.C.; Huat, J.; Agbossou, E.K.; Saint-Macary, H.; Glèlè Kakai, R. Vegetable Contamination by the Fecal Bacteria of Poultry Manure: Case Study of Gardening Sites in Southern Benin. International Journal of Food Science 2016, 2016, e4767453, doi:10.1155/2016/4767453.

40. Graham, J.P.; Evans, S.L.; Price, L.B.; Silbergeld, E.K. Fate of Antimicrobial-Resistant Enterococci and Staphylococci and Resistance Determinants in Stored Poultry Litter. Environmental Research 2009, 109, 682–689, doi:10.1016/j.envres.2009.05.005.

41. Thomas, K.V.; Bijlsma, L.; Castiglioni, S.; Covaci, A.; Emke, E.; Grabic, R.; Hernández, F.; Karolak, S.; Kasprzyk-Hordern, B.; Lindberg, R.H.; et al. Comparing Illicit Drug Use in 19 European Cities through Sewage Analysis. Science of The Total Environment 2012, 432, 432–439, doi:10.1016/j.scitotenv.2012.06.069.

42. Gurmessa, B.; Ashworth, A.J.; Yang, Y.; Savin, M.; Moore, P.A.; Ricke, S.C.; Corti, G.; Pedretti, E.F.; Cocco, S. Variations in Bacterial Community Structure and Antimicrobial Resistance Gene Abundance in Cattle Manure and Poultry Litter. Environmental Research 2021, 197, 111011, doi:10.1016/j.envres.2021.111011.

43. Lauber, C.L.; Hamady, M.; Knight, R.; Fierer, N. Pyrosequencing-Based Assessment of Soil PH as a Predictor of Soil Bacterial Community Structure at the Continental Scale. Applied and Environmental Microbiology 2009, 75, 5111–5120, doi:10.1128/AEM.00335-09.

44. Borda-Molina, D.; Vital, M.; Sommerfeld, V.; Rodehutscord, M.; Camarinha-Silva, A. Insights into Broilers’ Gut Microbiota Fed with Phosphorus, Calcium, and Phytase Supplemented Diets. Frontiers in Microbiology 2016, 7.

45. Rinninella, E.; Raoul, P.; Cintoni, M.; Franceschi, F.; Miggiano, G.A.D.; Gasbarrini, A.; Mele, M.C. What Is the Healthy Gut Microbiota Composition? A Changing Ecosystem across Age, Environment, Diet, and Diseases. Microorganisms 2019, 7, E14, doi:10.3390/microorganisms7010014.

46. Zhang, Y.; Gu, A.Z.; Cen, T.; Li, X.; He, M.; Li, D.; Chen, J. Sub-Inhibitory Concentrations of Heavy Metals Facilitate the Horizontal Transfer of Plasmid-Mediated Antibiotic Resistance Genes in Water Environment. Environ. Pollut. 2018, 237, 74–82, doi:10.1016/j.envpol.2018.01.032.

47. Peng, S.; Wang, Y.; Chen, R.; Lin, X. Chicken Manure and Mushroom Residues Affect Soil Bacterial Community Structure but Not the Bacterial Resistome When Applied at the Same Rate of Nitrogen for 3 Years. Frontiers in Microbiology 2021, 12.

48. Ziganshin, A.M.; Liebetrau, J.; Pröter, J.; Kleinsteuber, S. Microbial Community Structure and Dynamics during Anaerobic Digestion of Various Agricultural Waste Materials. Appl Microbiol Biotechnol 2013, 97, 5161–5174, doi:10.1007/s00253-013-4867-0.

49. Deng, W.; Zhang, A.; Chen, S.; He, X.; Jin, L.; Yu, X.; Yang, S.; Li, B.; Fan, L.; Ji, L.; et al. Heavy Metals, Antibiotics and Nutrients Affect the Bacterial Community and Resistance Genes in Chicken Manure Composting and Fertilized Soil. Journal of Environmental Management 2020, 257, 109980, doi:10.1016/j.jenvman.2019.109980.

50. Donaldson, G.P.; Lee, S.M.; Mazmanian, S.K. Gut Biogeography of the Bacterial Microbiota. Nat Rev Microbiol 2016, 14, 20–32, doi:10.1038/nrmicro3552.

51. Shafey, T.M.; McDonald, M.W.; Dingle, J.G. Effects of Dietary Calcium and Available Phosphorus Concentration on Digesta PH and on the Availability of Calcium, Iron, Magnesium and Zinc from the Intestinal Contents of Meat Chickens. British Poultry Science 1991, 32, 185–194, doi:10.1080/00071669108417339.

52. Ptak, A.; Bedford, M.R.; Swiatkiewicz, S.; Żyła, K.; Józefiak, D. Phytase Modulates Ileal Microbiota and Enhances Growth Performance of the Broiler Chickens. PLOS ONE 2015, 10, e0119770, doi:10.1371/journal.pone.0119770.

53. Witzig, M.; Silva, A.C. da; Green-Engert, R.; Hoelzle, K.; Zeller, E.; Seifert, J.; Hoelzle, L.E.; Rodehutscord, M. Spatial Variation of the Gut Microbiota in Broiler Chickens as Affected by Dietary Available Phosphorus and Assessed by T-RFLP Analysis and 454 Pyrosequencing. PLOS ONE 2015, 10, e0143442, doi:10.1371/journal.pone.0143442.

54. Peng, S.; Feng, Y.; Wang, Y.; Guo, X.; Chu, H.; Lin, X. Prevalence of Antibiotic Resistance Genes in Soils after Continually Applied with Different Manure for 30 Years. Journal of Hazardous Materials 2017, 340, 16–25, doi:10.1016/j.jhazmat.2017.06.059.

55. Qian, X.; Gu, J.; Sun, W.; Wang, X.-J.; Su, J.-Q.; Stedfeld, R. Diversity, Abundance, and Persistence of Antibiotic Resistance Genes in Various Types of Animal Manure Following Industrial Composting. J. Hazard. Mater. 2018, 344, 716–722, doi:10.1016/j.jhazmat.2017.11.020.

56. Xu, Y.; Li, H.; Shi, R.; Lv, J.; Li, B.; Yang, F.; Zheng, X.; Xu, J. Antibiotic Resistance Genes in Different Animal Manures and Their Derived Organic Fertilizer. Environmental Sciences Europe 2020, 32, 102, doi:10.1186/s12302-020-00381-y.

57. Wang, Y.; Hu, Y.; Liu, F.; Cao, J.; Lv, N.; Zhu, B.; Zhang, G.; Gao, G.F. Integrated Metagenomic and Metatranscriptomic Profiling Reveals Differentially Expressed Resistomes in Human, Chicken, and Pig Gut Microbiomes. Environment International 2020, 138, 105649, doi:10.1016/j.envint.2020.105649.

58. Zhao, L.; Dong, Y.H.; Wang, H. Residues of Veterinary Antibiotics in Manures from Feedlot Livestock in Eight Provinces of China. Sci Total Environ 2010, 408, 1069–1075, doi:10.1016/j.scitotenv.2009.11.014.

59. Antibiotics and Antimicrobial Resistance Genes: Environmental Occurrence and Treatment Technologies; Hashmi, M.Z., Ed.; Emerging Contaminants and Associated Treatment Technologies; Springer International Publishing, 2020; ISBN 978-3-030-40421-5.

60. Chen, J.; Fluharty, F.L.; St-Pierre, N.; Morrison, M.; Yu, Z. Technical Note: Occurrence in Fecal Microbiota of Genes Conferring Resistance to Both Macrolide-Lincosamide-Streptogramin B and Tetracyclines Concomitant with Feeding of Beef Cattle with Tylosin. J. Anim. Sci. 2008, 86, 2385–2391, doi:10.2527/jas.2007-0705.

61. Lee, S.; Mir, R.A.; Park, S.H.; Kim, D.; Kim, H.-Y.; Boughton, R.K.; Morris, J.G.; Jeong, K.C. Prevalence of Extended-Spectrum β-Lactamases in the Local Farm Environment and Livestock: Challenges to Mitigate Antimicrobial Resistance. Critical Reviews in Microbiology 2020, 46, 1–14, doi:10.1080/1040841X.2020.1715339.

62. Binh, C.T.T.; Heuer, H.; Kaupenjohann, M.; Smalla, K. Piggery Manure Used for Soil Fertilization Is a Reservoir for Transferable Antibiotic Resistance Plasmids. FEMS Microbiology Ecology 2008, 66, 25–37, doi:10.1111/j.1574-6941.2008.00526.x.

63. Moodley, A.; Guardabassi, L. Transmission of IncN Plasmids Carrying BlaCTX-M-1 between Commensal Escherichia Coli in Pigs and Farm Workers. Antimicrobial Agents and Chemotherapy 2009, 53, 1709–1711, doi:10.1128/AAC.01014-08.

64. Xiong, W.; Wang, Y.; Sun, Y.; Ma, L.; Zeng, Q.; Jiang, X.; Li, A.; Zeng, Z.; Zhang, T. Antibiotic-Mediated Changes in the Fecal Microbiome of Broiler Chickens Define the Incidence of Antibiotic Resistance Genes. Microbiome 2018, 6, 34, doi:10.1186/s40168-018-0419-2.

65. Guo, T.; Lou, C.; Zhai, W.; Tang, X.; Hashmi, M.Z.; Murtaza, R.; Li, Y.; Liu, X.; Xu, J. Increased Occurrence of Heavy Metals, Antibiotics and Resistance Genes in Surface Soil after Long-Term Application of Manure. Science of The Total Environment 2018, 635, 995–1003, doi:10.1016/j.scitotenv.2018.04.194.

66. Mahmoud, M.A.M.; Abdel-Mohsein, H.S. Hysterical Tetracycline in Intensive Poultry Farms Accountable for Substantial Gene Resistance, Health and Ecological Risk in Egypt-Manure and Fish. Environmental Pollution 2019, 255, 113039, doi:10.1016/j.envpol.2019.113039.

67. Yoshizawa, N.; Usui, M.; Fukuda, A.; Asai, T.; Higuchi, H.; Okamoto, E.; Seki, K.; Takada, H.; Tamura, Y. Manure Compost Is a Potential Source of Tetracycline-Resistant Escherichia Coli and Tetracycline Resistance Genes in Japanese Farms. Antibiotics 2020, 9, 76, doi:10.3390/antibiotics9020076.

68. Agersø, Y.; Pedersen, A.G.; Aarestrup, F.M. Identification of Tn5397-like and Tn916-like Transposons and Diversity of the Tetracycline Resistance Gene Tet(M) in Enterococci from Humans, Pigs and Poultry. Journal of Antimicrobial Chemotherapy 2006, 57, 832–839, doi:10.1093/jac/dkl069.

69. Leclercq, S.O.; Wang, C.; Zhu, Y.; Wu, H.; Du, X.; Liu, Z.; Feng, J. Diversity of the Tetracycline Mobilome within a Chinese Pig Manure Sample. Applied and Environmental Microbiology 2016, 82, 6454–6462, doi:10.1128/AEM.01754-16.

70. Wang, F.-H.; Qiao, M.; Su, J.-Q.; Chen, Z.; Zhou, X.; Zhu, Y.-G. High Throughput Profiling of Antibiotic Resistance Genes in Urban Park Soils with Reclaimed Water Irrigation. Environ Sci Technol 2014, 48, 9079–9085, doi:10.1021/es502615e.

71. Domínguez, M.; Miranda, C.D.; Fuentes, O.; de la Fuente, M.; Godoy, F.A.; Bello-Toledo, H.; González-Rocha, G. Occurrence of Transferable Integrons and Sul and Dfr Genes Among Sulfonamide-and/or Trimethoprim-Resistant Bacteria Isolated From Chilean Salmonid Farms. Frontiers in Microbiology 2019, 10.

72. Møller, A.K.; Barkay, T.; Hansen, M.A.; Norman, A.; Hansen, L.H.; Sørensen, S.J.; Boyd, E.S.; Kroer, N. Mercuric Reductase Genes (MerA) and Mercury Resistance Plasmids in High Arctic Snow, Freshwater and Sea-Ice Brine. FEMS Microbiology Ecology 2014, 87, 52–63, doi:10.1111/1574-6941.12189.

73. Popowska, M.; Krawczyk-Balska, A. Broad-Host-Range IncP-1 Plasmids and Their Resistance Potential. Frontiers in Microbiology 2013, 4.

74. Zhou, Y.; Awasthi, S.K.; Liu, T.; Verma, S.; Zhang, Z.; Pandey, A.; Varjani, S.; Li, R.; Taherzadeh, M.J.; Awasthi, M.K. Patterns of Heavy Metal Resistant Bacterial Community Succession Influenced by Biochar Amendment during Poultry Manure Composting. Journal of Hazardous Materials 2021, 420, 126562, doi:10.1016/j.jhazmat.2021.126562.

75. Qiu, X.; Zhou, G.; Wang, H.; Wu, X. The Behavior of Antibiotic-Resistance Genes and Their Relationships with the Bacterial Community and Heavy Metals during Sewage Sludge Composting. Ecotoxicology and Environmental Safety 2021, 216, 112190, doi:10.1016/j.ecoenv.2021.112190.

76. Liao, H.; Zhao, Q.; Cui, P.; Chen, Z.; Yu, Z.; Geisen, S.; Friman, V.-P.; Zhou, S. Efficient Reduction of Antibiotic Residues and Associated Resistance Genes in Tylosin Antibiotic Fermentation Waste Using Hyperthermophilic Composting. Environment International 2019, 133, 105203, doi:10.1016/j.envint.2019.105203.

77. Biemer, J.J. Antimicrobial Susceptibility Testing by the Kirby-Bauer Disc Diffusion Method. Ann Clin Lab Sci 1973, 3, 135–140.

78. Ashfaq, M.Y.; Da’na, D.A.; Al-Ghouti, M.A. Application of MALDI-TOF MS for Identification of Environmental Bacteria: A Review. Journal of Environmental Management 2022, 305, 114359, doi:10.1016/j.jenvman.2021.114359.

79. Hogan, C.A.; Watz, N.; Budvytiene, I.; Banaei, N. Rapid Antimicrobial Susceptibility Testing by VITEK®2 Directly from Blood Cultures in Patients with Gram-Negative Rod Bacteremia. Diagn Microbiol Infect Dis 2019, 94, 116–121, doi:10.1016/j.diagmicrobio.2019.01.001.

80. Bolyen, E.; Rideout, J.R.; Dillon, M.R.; Bokulich, N.A.; Abnet, C.C.; Al-Ghalith, G.A.; Alexander, H.; Alm, E.J.; Arumugam, M.; Asnicar, F.; et al. Reproducible, Interactive, Scalable and Extensible Microbiome Data Science Using QIIME 2. Nat Biotechnol 2019, 37, 852–857, doi:10.1038/s41587-019-0209-9.

81. Chong, J.; Liu, P.; Zhou, G.; Xia, J. Using MicrobiomeAnalyst for Comprehensive Statistical, Functional, and Meta-Analysis of Microbiome Data. Nat Protoc 2020, 15, 799–821, doi:10.1038/s41596-019-0264-1.

82. Chen, Q.; An, X.; Li, H.; Su, J.; Ma, Y.; Zhu, Y.-G. Long-Term Field Application of Sewage Sludge Increases the Abundance of Antibiotic Resistance Genes in Soil. Environment International 2016, 92–93, 1–10, doi:10.1016/j.envint.2016.03.026.

