## Supplementary figures and images for "A comprehensive study of the microbiome and resistome of chicken waste from intensive farms"

### Figure S1

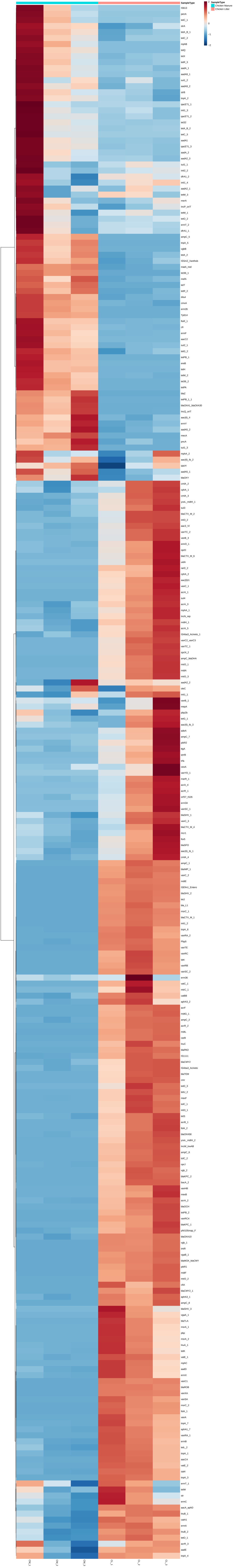
