## Supplementary material for "A comprehensive study of the microbiome and resistome of chicken waste from intensive farms": Table S1-S3

**Table S1.** Antibiotic resistance [%] of isolated strains form two types of chicken wastes Chicken Litter (CL) and Chicken Manure (CM)

| **Antibiotic (n=total number of strains tested)** | **Sensitive strains (%)** | **Intermediate strains (%)** | **Resistant strains (%)** |
|---|---|---|---|
| Amoxicillin/Clavulanic acid (n=57) | 29.8% | 7% | 63.2% |
| Piperacillin/Tazobactam (n=57) | 59.7% | 23.6% | 16.7% |
| Cefuroxime (n=57) | 56.1% | 7.1% | 36.8% |
| Ceftazidime (n=57) | 45.6% | 1.8% | 52.6% |
| Imipenem (n=72) | 58.3% | 34.7% | 7% |
| Meropenem (n=72) | 94.4% | 2.8% | 2.8% |
| Gentamicin (n=91) | 49.5% | - | 50.5% |
| Amikacin (n=72) | 80.5% | 1.4% | 18.1% |
| Tobramycin (n=72) | 45.8% | - | 54.2% |
| Cefotaxime (n=57) | 50% | 1.7% | 48.3% |
| Ciprofloxacine (n=91) | 25.3% | 13.2% | 61.5% |
| Cefepime (n=57) | 94.8% | 3.5% | 1.7% |
| Cefuroxime-axetil (n=57) | 45.6% | - | 54.4% |
| Tigecycline (n=77) | 62.3% | 1.3% | 36.4% |
| Trimetoprim/Sulfametoxazol (n=90) | 72.2% | 2.2% | 25.6% |
| Levofloxacin (n=34) | 14.7% | 11.8% | 73.5% |
| Erythromycin (n=19)* | 42.1% | - | 47.9% |
| Tetracycline (n=19)* | - | - | 100% |
| Clindamycin (n=19)* | 10.5% | 10.5% | 79% |

*only *Staphylococcus luteus* strains

**Table S2**. Selective agar screening for AMR pathogens

| Bacteria species | Screening selective agar + antibiotic (Concentration) using the EUCAST breakpoints | Selecting for |
|---|---|---|
| *Klebsiella pneumoniae* | Simmons Citrate Agar + inositol Imipenem (16 mg/L) | Carbapenem resistance |
| *Klebsiella pneumoniae* | Simmons Citrate Agar + inositol Cefotaxime (4 mg/L) | Extended-spectrum beta-lactamase |
| *Escherichia coli* | Eosin Methylene Blue Agar Imipenem (16 mg/L) | Carbapenem resistance |
| | MacConkey Agar Imipenem (16 mg/L) | |
| *Escherichia coli* | Eosin Methylene Blue Agar Cefotaxime (4 mg/L) | Extended-spectrum beta-lactamase |
| | MacConkey Agar Cefotaxime (4 mg/L) | |
| *Enterococcus sp.* | Brilliance VRE Agar | Vancomycin resistance |
| *Acinetobacter baumannii* | CHROMagar Acinetobacter Imipenem (16 mg/L) | Carbapenem resistance |
| *Pseudomonas aeruginosa* | Cetrimide Agar Imipenem (16 mg/L) | Carbapenem resistance |
| *Staphylococcus aureus* | Mannitol salt Agar Oxacillin (4 mg/L) | Methicillin resistance |
| *Staphylococcus aureus* | Mannitol Salt Agar Vancomycin (4 mg/L) | Vancomycin resistance |

**Table S3.** Antimicrobial susceptibility discs used for the isolated strains' phenotypic characterization

| Selective medium | Antibiotics (abbreviation, concentration) |
|---|---|
| Simmons Citrate Agar | IMP, 10 µg; CIP, 5 µg; CTX, 5 µg; CN, 10 µg |
| Eosin Methylene Blue Agar | IMP (10 µg), CIP (5 µg), CTX (5 µg), CN (10 µg) |
| MacConkey Agar | IMP (10 µg), CIP (5 µg), CTX (5 µg), CN (10 µg) |
| Brilliance VRE agar | IMP (10 µg), CIP (5 µg), VAN (30 µg), QD (15 µg) |
| Cetrimide Agar | IMP (10 µg), CIP (5 µg), CTX (5 µg), FEP (30 µg) |
| Mannitol Salt Agar | OX (1 µg), TE (30 µg), VAN (30 µg), ER (15 µg) |

**IMP** - imipenem, **CIP** – ciprofloxacin, **CTX** – cefotaxime, **CN** – gentamicin, **VAN** – vancomycin, **QD** - quinupristin/dalfopristin, **FEP** – cefepime, **OX** – oxacillin, **TE** – tetracycline, **ER** - erythromycin
